# *Bacopa monnieri (L.)* Wettst-derived nanovesicles are enriched with bioactive cargo and exhibit anti-neuroblastoma activity

**DOI:** 10.64898/2026.01.17.700097

**Authors:** Ekkaphot Khongkla, Panitch Boonsnongcheep, Pipob Suwanchaikasem, Kornkanok Promthep, Monruedee Srisaisup, Theptharin Charuraksa, Pannaphan Makarathut, Banthit Chetsawang

## Abstract

Edible plant-derived nanovesicles (PDNVs) have emerged as promising nanotherapeutic strategies for various diseases, including cancer. The biochemical composition and functional properties of PDNVs vary considerably on the basis of their botanical source. *Bacopa monnieri* (L.) Wettst is a medicinal plant renowned for its rich phytochemical profile, yet the isolation and biological activities of *B. monnieri* -derived nanovesicles (BMNVs) remain unexplored. We report, for the first time, the isolation, molecular cargo profiling, and *in vitro* functional evaluation of BMNVs against neuroblastoma cells. The isolated BMNVs displayed a characteristic bilayer morphology with an average particle size of ∼112 nm. Mass spectrometry-based metabolite analysis revealed an enrichment of triterpenoids and triterpene saponins, whereas protein cargo analysis revealed superoxide dismutase, which is correlated with their intrinsic free radical scavenging activity. *In vitro* assays demonstrated that BMNVs significantly suppress neuroblastoma cell growth and induce morphological alterations. Confocal three-dimensional reconstruction confirmed the cellular internalization of the BMNVs, revealing a distinct perinuclear distribution. This study provides the first evidence of the use of BMNVs as bioactive carriers, highlighting their potential as novel nanotherapeutic agents and establishing *B. monnieri* as a valuable natural resource for the development of bioactive plant-derived nanovesicles for nanomedicine.

**Highlights:**

- First isolation and biophysical characterization of *Bacopa monnieri*-derived nanovesicles (BMNVs).
- BMNVs have diverse metabolite profiles and are notably enriched in triterpenoids and triterpene saponins.
- The superoxide dismutase (SOD) identified within BMNVs confers intrinsic free radical-scavenging activity.
- BMNVs exhibit therapeutic potential as anti-neuroblastoma agents.
- These edible plant-derived nanovesicles offer a versatile, biogenic platform to explore for further development in diverse therapeutic and nutraceutical applications.

## 1. Introduction

Neuroblastoma is an aggressive pediatric embryonal tumor originating from sympathoadrenal neuroblasts and remains a leading cause of childhood cancer mortality, often presenting at advanced stages with high relapse rates and substantial long-term toxicity from multimodal therapy [1,2]. These limitations have spurred growing interest in more biocompatible nanomedicine-based strategies for targeting neuroblastoma [3].

Plant-derived nanovesicles (PDNVs) are nanosized lipid bilayer vesicles isolated from botanical resources and have emerged as promising innovations in health-related research [4–7]. PDNVs act as messengers that play pivotal roles in intercellular communication by transferring bioactive cargo, such as proteins, lipids, RNAs, and secondary metabolites, which are capable of modulating the biological functions and signaling pathways of recipient cells. Notably, increasing evidence indicates that PDNVs participate in cross-kingdom interactions, whereby vesicles released from plants are internalized by mammalian cells, highlighting the potential of PDNVs as naturally derived regulators of mammalian cellular functions. Additionally, their inherent advantages, including excellent biocompatibility, low immunogenicity, abundant availability, ease of large-scale isolation, and sustainable production, support their development as nanotherapeutic platforms for a wide range of medical and biological applications.

A growing body of evidence indicates that PDNVs exert potent anticancer effects. For example, Ginger-derived exosome-like nanoparticles have been shown to induce G2/M phase cell cycle arrest in human breast adenocarcinoma cells [8]. In addition, extracellular vesicles derived from *Salvia dominica* hairy roots exhibit marked cytotoxicity in cancer cells, with efficacy comparable to that of gemcitabine, a chemotherapeutic agent that targets the DNA damage response [9].

These nanovesicles are enriched with diverse bioactive molecules capable of modulating multiple signaling pathways involved in cancer cell proliferation and survival. In neuroblastoma models, extracellular vesicles derived from Coffea arabica beans significantly reduced SH-SY5Y cell viability[10]. However, current evidence in neuroblastoma is restricted to a small number of plant species. To further advance the development of PDNVs as nanotherapeutics for future neuroblastoma therapy, the identification and characterization of nanovesicles derived from additional plant sources, particularly from medicinal and edible plants that harbor intrinsically bioactive cargo, are needed.

*Bacopa monnieri* (L.) Wettst is a medicinal creeping perennial plant native to India and Southeast Asia, including Thailand, and belongs to the Plantaginaceae family. The plant is characterized by small oblong leaves and white to pale purple flowers and has been extensively utilized in traditional Ayurvedic medicine[11]. *B. monnieri* (BM) is widely recognized for its therapeutic benefits, particularly in cognitive enhancement, as well as its applications in the management of insomnia, epilepsy, and anxiety-related disorders[12–15]. Preclinical *in vivo* studies have demonstrated the absence of acute and subchronic toxicity even at relatively high doses of BM extracts, supporting its favorable safety profile [16,17]. Moreover, to date, there have been no reported cases of adverse effects or toxicity associated with BM consumption in humans, further reinforcing its status as a safe medicinal herb[18]. Consequently, BM is generally regarded as an edible and pharmacologically valuable plant, making it an attractive candidate for the development of novel therapeutic platforms.

Accumulating evidence has demonstrated that BM extracts exhibit anticancer activities [19–21]. Triterpenoid saponins serve as the primary bioactive constituents underlying these pharmacological effects. Among these compounds, bacosides, especially bacoside A, are the most extensively characterized and are widely recognized for their broad spectrum of biological efficacy [22–24]. Previous studies have reported the *in vitro* bioactivity of bacoside A against several cancer cell lines [25], including the induction of cell cycle arrest in glioblastoma U-87 MG cells [26]. Collectively, these findings highlight the tumor-suppressing potential of BM-derived bioactive molecules. Despite these promising pharmacological properties, the clinical and translational application of bacosides is limited by their relatively low bioavailability and stability, which restrict their effective therapeutic efficacy. This limitation underscores the need for alternative strategies that enable more efficient plant utilization and facilitate the effective translation of its therapeutic benefits. One emerging approach involves the exploitation of BM-derived nanovesicles (BMNVs). To date, there is no experimental evidence supporting the isolation, characterization, and biological functions of BMNVs, representing a significant knowledge gap and hindering their translational potential.

This study reports, for the first time, the successful isolation and characterization of BMNVs. We identified their molecular composition and evaluated their biological activities, revealing a cargo enriched with triterpenoids, triterpene saponins, and superoxide dismutase. The potent growth-suppressive effects and efficient cellular internalization in SH-SY5Y neuroblastoma cells underscore the bioactivity of BMNVs, supporting their prospective development as novel nanotherapeutic and delivery agents.

## 2. Materials and methods

### 2.1 Cell culture

The human neuroblastoma SH-SY5Y (cat.no. iCell-h187) cell line was purchased from iCell Bioscience, China. The cells were maintained in Dulbecco’s modified Eagle’s medium (DMEM): nutrient mixture F-12 (F12) (Gibco, USA) supplemented with 10% (v/v) fetal bovine serum (FBS) (HyClone, USA) and 1% (v/v) antibiotic-antimycotic 100X (Gibco, USA). The human kidney HEK293 cell line obtained from the American Type Culture Collection (ATCC; CRL-1573) was cultured in high-glucose DMEM (cat. no. D5648; Sigma Aldrich, USA) supplemented with 10% FBS and 1% antibiotics. These cells were grown in a humidified incubator at 37°C with 5% CO_2_ and 95% air (v/v). The cultured cells were regularly tested for mycoplasma contamination via a Mycoplasma PCR detection kit (Abcam, UK).

### 2.2 Isolation of BMNVs

Fresh and cleaned aerial parts of *B. monnieri* were purchased from local plant market and identified by Dr. Panitch Boonsnongcheep. The plants were homogenized via a blender. The resulting homogenate was subjected to sequential filtration through white filter papers to remove large fibrous materials. The filtrate was then centrifuged at 10,000 × g for 60 min at 4°C for two consecutive rounds to eliminate residual cellular debris. The resulting supernatants were collected and further centrifuged at 50,000 × g for 60 min at 4°C for two rounds. The supernatants were subsequently passed through a 0.45 μm filter and ultracentrifuged at 100,000 × g for 60 min at 4°C for two rounds to pellet the extracellular vesicles. To increase vesicle purity, the pellets were subjected to ultrafiltration via Macrosep centrifugal filters (MAP100C37, Cytiva, USA) to remove low-molecular-weight components and soluble contaminants. The retained fraction was then centrifuged at 3,000 × g for 15 min for three consecutive rounds to recover *B. monnieri*–derived nanovesicles (BMNVs). The final suspension was filtered through a 0.45 μm membrane and stored at −30°C for short-term use and −80°C for long-term storage.

### 2.3 Nanoparticle tracking analysis (NTA)

The size distribution and concentration of the BMNVs were analyzed via NanoSight Pro (Malvern Panalytical, UK). The samples (200 μL) were injected with a sterile syringe. For each measurement, three videos were captured. The measurements were performed at room temperature.

### 2.4 Cryogenic electron microscopy (cryo-EM)

To visualize the morphology of the isolated BMNVs. The samples were submitted to the Research Frontier Facility (FRF), Mahidol University. An aliquot of each sample was applied to a plasma-cleaned copper grid (C-flat R 1.2/1.3, 300 mesh, Electron Microscopy Sciences) and prepared via an automated grid plunger (Vitrobot Mark IV, Thermo Fisher Scientific) with the environmental chamber set at 100% humidity and 4°C. The grid was blotted for 4.5 s and vitrified in liquid ethane. The frozen hydrated exosomes were imaged via a cryogenic electron microscope (Glacios, Thermo Fisher Scientific) equipped with a Falcon 3EC. Cryo-EM imaging was performed at 73,000x and 120,000x magnification at 200 kV.

### 2.5 Zeta potential measurement

Zeta potential analysis was performed to determine the surface charge and colloidal stability of the isolated nanovesicles[27]. The samples 50 μL were diluted in sterile distilled water (1 mL) and applied to the nanoPartica SZ-100V2 nanoparticle analyzer (Horiba). Zeta potential analysis was carried out using standard setting (dispersion medium viscosity = 0.896 mPa·s, temperature = 25 °C). The data were analyzed by the Horiba SZ-100 software version 2.44.

### 2.6 Metabolite identification

#### 2.6.1 Metabolite extraction

Metabolite profiling of lyophilized BMNVs was performed via a liquid chromatography–quadrupole time‒of‒flight mass spectrometry (LC–QTOF–MS) platform. Briefly, samples were extracted with 200 µL of 70% methanol containing sulfadimethoxine (50 ng/mL) as an internal standard to enable subsequent signal normalization. The mixture was thoroughly vortexed and subjected to ultrasonic extraction for 10 min at room temperature to facilitate metabolite solubilization. Following extraction, the samples were centrifuged at 14,000 rpm for 10 min, and the clarified supernatants were carefully transferred into liquid chromatography–mass spectrometry (LC–MS) vials for instrumental analysis.

#### 2.6.2 Sample acquisition

Chromatographic separation was conducted on an Agilent LC system equipped with a Poroshell 120 EC-C18 column (2.1 × 100 mm, 2.7 µm particle size) maintained at 50°C. The injection volume was set to 10 µL, and the flow rate was maintained at 0.4 mL/min throughout the analysis. Mobile phase A consisted of water containing 0.1% formic acid, while mobile phase B consisted of acetonitrile supplemented with 0.1% formic acid. The gradient elution program was optimized for metabolite separation and followed the conditions in Table 1, with an initial high aqueous phase, a gradual increase in organic solvent, and a final re-equilibration step to restore the initial conditions.

**Table 1.** Gradient elution scheme for liquid chromatography for metabolite analysis.

| Time (min) | % A | % B |
| --- | --- | --- |
| 0.0 | 100 | 0 |
| 0.5 | 100 | 0 |
| 10.5 | 45 | 55 |
| 12.5 | 25 | 75 |
| 14.0 | 0 | 100 |
| 17.0 | 0 | 100 |
| 17.5 | 100 | 0 |
| 20.0 | 100 | 0 |

Mass spectrometric detection was performed via an Agilent LC-QTOF 6545XT operated in high-resolution acquisition mode under both positive and negative electrospray ionization (ESI) conditions. The drying gas temperature was set to 325°C with a flow rate of 13 L/h, while the sheath gas temperature and flow rate were maintained at 275°C and 12 L/h, respectively. The nebulizer pressure was set at 45 psi. Capillary voltages were configured at 4000 V for positive mode and 3000 V for negative mode. The isotope width was set to 1.3 m/z. Full-scan MS (MS1) data were acquired over a mass range of 40–1700 m/z, and MS/MS (MS2) spectra were collected from 25–1000 m/z via data-dependent acquisition. The collision energy was fixed at 20 eV in positive mode and 10 eV in negative mode. The acquisition rate was set to 3.35 spectra per second, with a maximum of 10 precursors selected per cycle and a precursor ion intensity threshold of 5000 counts. Continuous mass axis calibration was achieved via reference ions at m/z 121.0509 and 922.0098 in positive mode and m/z 112.9856 and 1033.9881 in negative mode.

#### 2.6.3 Data processing and analysis

The raw data were processed via MS-DIAL software version 5.3. Feature alignment was performed against the exosome sample dataset, and signal intensities were normalized to the sulfadimethoxine internal standard. Background and instrumental artifacts were minimized by excluding specified reference and contaminant ions, including m/z 121.0509 and 922.0098 in positive mode and m/z 112.9856, 119.0363, 966.0007, and 1033.9881 in negative mode. Metabolite annotation considered M+H adducts in positive mode and M−H adducts in negative mode. Metabolite identification was carried out by matching accurate mass, retention time, and MS/MS fragmentation patterns against multiple spectral libraries, including the MS-DIAL ESI (+/−) MS/MS database from authentic standards, the Fiehn/Vaniya natural product library, and the BMDMS-NP database[28]. The identification criteria included a retention time window of 0.2–18 min, a sample-to-blank ratio of at least 50, an identification score ≥ 0.7, and a mass error tolerance within ±20 ppm. Using these stringent criteria, a total of 120 metabolic features were detected in positive ionization mode, and 43 features were detected in negative ionization mode. Detailed information on the annotated metabolites is provided in Supplementary File 1.

### 2.7 Protein identification

#### 2.7.1 Sample preparation

The lyophilized BMNVs were resuspended in 60 µl of ammonium bicarbonate (50 mM) buffer. The sample was centrifuged at 14,000 rpm for 10 min, and the supernatant was collected for analysis. The protein mixture (40 µl) was reduced with 10 mM dithiothreitol (DTT) at 65°C for 30 min. Iodoacetamide (25 mM) was added for alkylation, and the sample was incubated at room temperature (RT) in the dark for 20 min. For protein digestion, 2.5 µl of 0.1 mg/mL trypsin was added, and the sample was incubated at 37°C overnight. The digestion reaction was terminated by the addition of 10% formic acid. The samples were subsequently centrifuged at 14,000 rpm for 10 min to collect the supernatant, which was subsequently transferred to an LC‒MS vial for subsequent LC‒MS analysis.

#### 2.7.2 Sample acquisition

A volume of 20 µL of the prepared sample was injected into the LC system for analysis. Chromatographic separation was performed on an Agilent Peptide Map column (2.1 × 150 mm, 2.7 µm particle size) maintained at a constant temperature of 60°C to ensure optimal peak shape and reproducibility. The mobile phases consisted of mobile phase A: water containing 0.1% (v/v) formic acid and mobile phase B: acetonitrile containing 0.1% (v/v) formic acid. The peptides were eluted at a constant flow rate of 0.4 mL/min via a linear multistep gradient over a total run time of 85 minutes. The gradient program is shown in Table 2.

**Table 2.** Gradient elution scheme for liquid chromatography for protein identification.

| Time (min) | %A | %B |
| --- | --- | --- |
| 0 | 100 | 0 |
| 2 | 100 | 0 |
| 35 | 80 | 20 |
| 55 | 70 | 30 |
| 65 | 50 | 50 |
| 70 | 10 | 90 |
| 75 | 10 | 90 |
| 80 | 100 | 0 |
| 85 | 100 | 0 |

Mass spectrometric analysis was carried out via an Agilent LC-QTOF 6545XT mass spectrometer equipped with an electrospray ionization (ESI) source operated in positive ion mode. The source parameters were optimized as follows: gas temperature, 325°C; drying gas flow, 13 L/h; nebulizer pressure, 35 psi; capillary voltage, 4000 V; and nozzle voltage, 500 V. Full-scan MS data were acquired over a mass range of m/z 100–1700, enabling comprehensive detection of peptide precursor ions. Data-dependent MS/MS acquisition was performed in centroid mode, with fragment ion spectra collected over an MS/MS mass range of m/z 25–1000. Collision energies were applied dynamically on the basis of the precursor charge state and m/z value. For +1- and +2-charged precursor ions, the collision energy was calculated via the equation CE = 3.1 × ((m/z)/100) + 1. For precursor ions with charge states ≥ +3, the collision energy was calculated as CE = 3.6 × ((m/z)/100) − 4.8. A continuous reference mass of m/z 922.0098 was used for internal mass calibration to ensure high mass accuracy throughout the acquisition.

#### 2.7.3 Data processing and protein identification

Raw data files acquired in Agilent proprietary (.d) format were converted to mzXML format via the MSConvert and OpenMS tools prior to database searching. Protein identification and label-free quantification were carried out via MaxQuant software (version 2.6.3). Spectral searches were performed against the UniProt protein databases, which were specifically restricted to two taxonomic levels: the Bacopa genus (taxonomy ID: 90645; 164 protein entries) and the Plantaginaceae family (taxonomy ID: 156152; 49,587 protein entries). Protein digestion was specified as trypsin, allowing for enzymatic cleavage at the C-terminal side of lysine and arginine residues. Carbamidomethylation of cysteine residues was defined as a fixed modification, whereas N-terminal acetylation and methionine oxidation were set as variable modifications. The minimum label-free quantification (LFQ) ratio was set to 1. The false discovery rate (FDR) threshold for both peptide-spectrum matches and protein identification was set at 0.1. To ensure high-confidence identification, proteins identified exclusively by site, reverse sequence hits, and known potential contaminants were systematically removed from the final dataset (Supplementary File 2).

### 2.8 DPPH radical scavenging assay

The antioxidant activity (AA) was monitored by the reduction reaction of the DPPH free radical (1,1-diphenyl-2-picrylhydrazyl). The samples were prepared with a 2-fold dilution from the highest concentration of BMNVs. The sample (100 µL) was mixed with 100 µL of DPPH solution (0.2 mM) freshly dissolved in methanol and incubated at room temperature in the dark for 30 min. Trolox (Sigma‒Aldrich, USA) was used as a standard antixodaint. The color change of the solution from an intense purple color to pale yellow was observed, and the absorbance values (A) were detected at 517 nm via a Biochrom EZ Read 800 microplate reader (Biochrom, UK).

The reduction in AB is proportional to the AA of the BMNVs. The AA was calculated according to the formula below, while the Trolox equivalent antioxidant capacity (TEAC) values were obtained from the linear regression equation of the Trolox calibration curve.

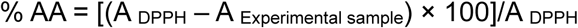

### 2.9 *In vitro* cytotoxicity

Human SH-SY5H or HEK293 cells were plated on 96-well plates at a density of 1 × 10^4^ cells/well for 24 h and exposed to different concentrations (particles/mL) of BMNV for 24 h. Doxorubicin (cat.no. 5927, Cell Signaling Technology, USA) at 5 μM was included as a positive control. The cytotoxicity was investigated via the 3-(4,5-dimethylthiazol-2-yl)-2,5-diphenyltetrazolium bromide (MTT) assay. In brief, MTT (Invitrogen, USA) at a final concentration of 0.5 mg/mL was added to each well for 4 h to form formazan. The crystals were solubilized with DMSO, and the absorbance was detected at 570 nm via a Biochrom EZ Read 800 microplate reader (Biochrom, UK).

### 2.10 Immunocytochemistry

The cells were fixed with 4% paraformaldehyde (PFA)/PBS for 15 min at room temperature and rinsed three times with PBS. The cells were then permeabilized with 0.2% Triton X-100 at room temperature and washed with PBS three times. To block nonspecific binding, the permeabilized cells were incubated with 1% bovine serum albumin (BSA)/PBS for 30 min at 37°C. The cells were costained with the following primary antibodies: anti-Ki67 (cat. no. ab15580, Abcam, UK) and anti-βIII-tubulin (cat. no. T8578, Sigma‒Aldrich, USA) on parafilm for 30 min at 37°C before being washed with PBS three times, each for more than 5 min. The cells were subsequently incubated with donkey anti-rabbit IgG (H+L) highly cross-adsorbed secondary antibody, Alexa Fluor 594 (cat. no. A21207, Invitrogen, USA) and donkey anti-mouse IgG-FITC (cat.no. sc-209, Santa Cruz Biotechnology, USA) for 30 min at 37°C. The final dilution was 1:200 for all the antibodies. The incubated cells were washed with PBS three times, washed with clean water, and mounted on slides using Fluoromount-G Mounting Medium with DAPI (cat. no. 00495952; Invitrogen, USA). Images were acquired via a Zeiss LSM 800 confocal laser scanning microscope.

### 2.11 Cellular uptake assay

SH-SY5Y cells were plated on a 12-well plate for 24 h at a density of 50,000 cells/well. The BMNVs were tagged with a PKH26 red fluorescent cell linker mini kit (MINI26, Sigma‒Aldrich, USA). The mixture was combined with 200 μl of the BMNVs, 4 μl of PKH26 dye, and 1 ml of Diluent C and incubated for 5 min at room temperature. The reaction was stopped by adding 10 mL of 1% BSA/PBS. The removal of excess free dye was performed via Macrosep advance centrifugal devices as described previously to collect the labeled BMNVs. To observe interanalization, the tagged BMNVs were resuspended in complete culture media and incubated with the cells for 16 h. The cells were fixed, permeabilized, and stained for βIII-tubulin as described in the immunocytochemistry section. The same samples were used for 2D and 3D reconstitution for visualization via a high-end Leica Stellaris STED microscope (Leica Microsystems CMS, Mannheim, Germany) via Z-Stack images and LasX 3D visualization software.

### 2.12 Statistical analysis

All statistical analyses were conducted using GraphPad Prism 8 (GraphPad Software Inc., USA). Data are expressed as the means ± standard deviations (SDs). Statistical significance was assessed using unpaired Student’s t tests for comparisons between two groups. For multiple group comparisons, one-way analysis of variance (ANOVA) followed by Tukey’s post hoc test was applied. For experiments involving two independent factors, two-way ANOVA followed by Tukey’s post hoc test was performed. Differences were considered statistically significant at P < 0.05.

## 3. Results

### 3.1 Nanovesicles derived from *B. monnieri* were characterized by their nanoscale size and the presence of a lipid bilayer membrane

*B. monnieri* is a small, nonaromatic, creeping herbaceous plant with a prostrate growth habit and numerous branching stems. The leaves are arranged oppositely in the stem (Figure 1). To isolate *B. monnieri*–derived nanovesicles (BMNVs), the aerial parts of the plants were harvested, and a combined ultracentrifugation and ultrafiltration approach was used. A diagram depicting the BMNV extraction and experimental workflow is shown in Figure 2.

**Figure 1.**
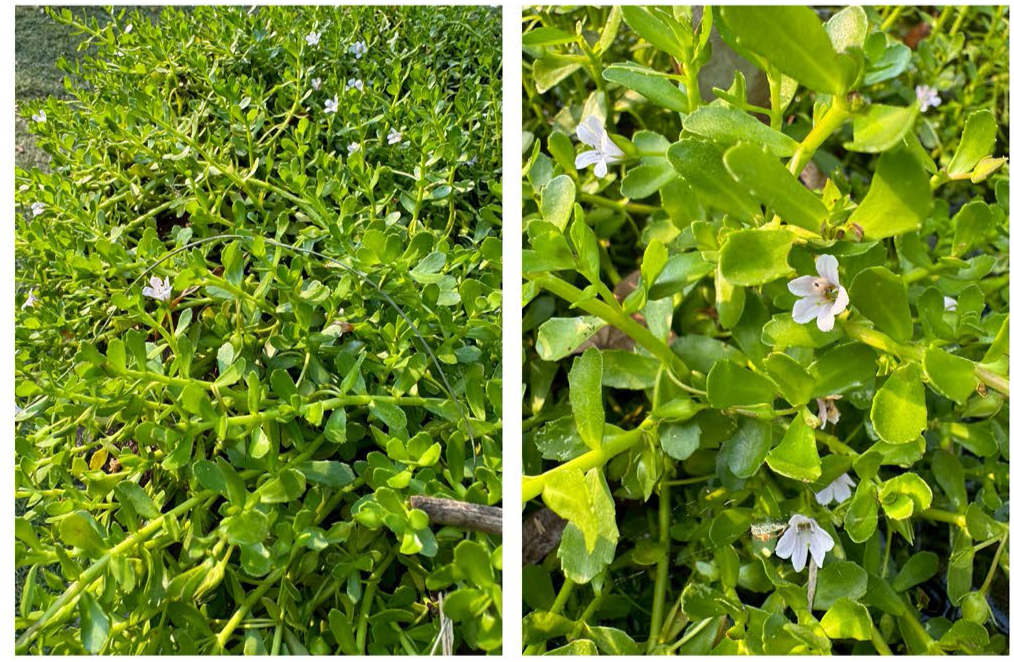
Representative images of the *Bacopa monnieri* (L.) Wettst plant used for this study.

**Figure 2.**
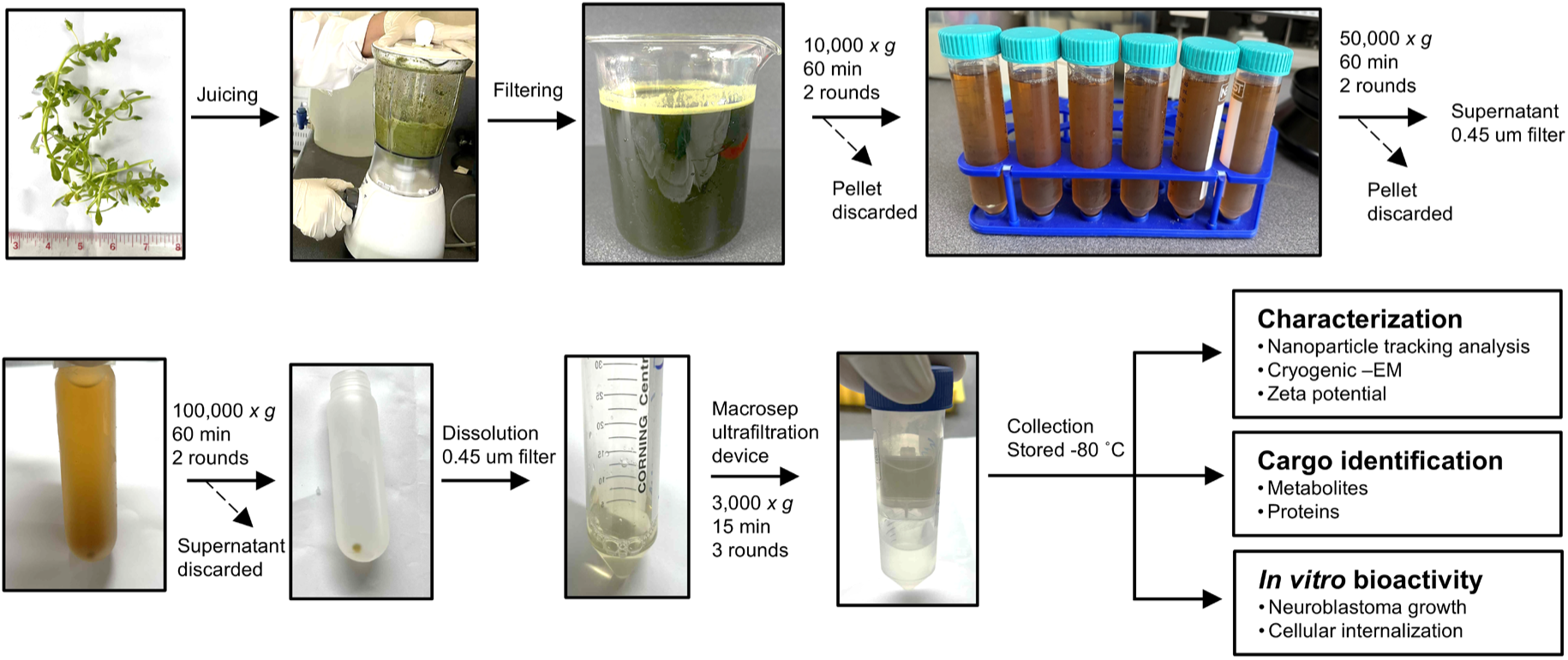
A diagram illustrating the steps of the isolation of BMNVs and downstream experiments, including characterization, composition identification, and *in vitro* bioactivity assessment.

The isolated BMMVs were characterized via biophysical methods following the Minimal Information for Studies of Extracellular Vesicles (MISEV) guidelines [29]. The concentration and size of the isolated BMNVs were measured via NTA (Figure 3a). The average sizes of the BMNVs were 112.5 nm (mode) and 173 nm (mean), confirming the nanoscale size of the particles. The lipid bilayer membrane measured by cryo-EM demonstrated the key characteristics of the vesicles (Figure 3b). Zeta potential average value was -27 ± 1.3 mV, indicating that the BMNVs have no tendency toward aggregated forms in suspension (Figure 3c). The stability of the BMNVs was also evaluated following six weeks of storage at −30°C. NTA revealed no statistically significant alterations in vesicle size or particle concentration compared with those of freshly isolated samples, indicating that the nanovesicles preserved their external physical properties during storage (Supplementary Figure 1).

**Figure 3.**
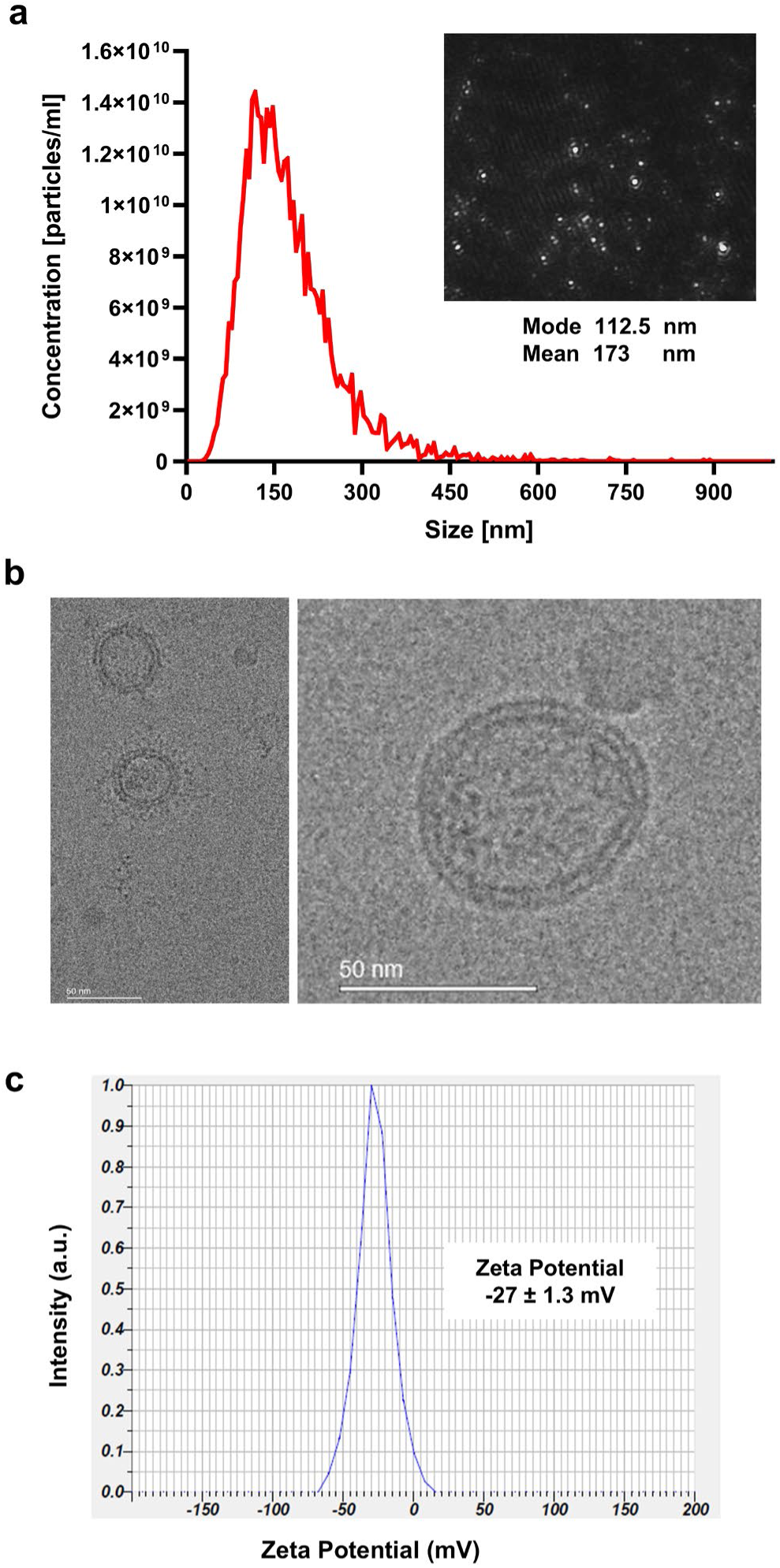
Characterization of BMNVS. **(a)** Nanoparticle tracking analysis of the isolated BMNVs to reveal the particle distribution (particle/mL) and average size (nm). **(b)** Representative cryo-EM images confirming the lipid bilayer structure of the BMNVs. The scale bar is 50 nm. **(c)** The zeta potential of isolated BMNVs was measured using a horiba nanoparticle analyzer sz-100. The measurements were conducted in triplicates.

### 3.2 BMNVs are enriched with diverse classes of metabolites, predominantly triterpenoids and triterpene saponins

To identify the bioactive compositions, present in the isolated BMNVs, mass spectrometry analysis was employed to profile and annotate the metabolites and proteins carried by the nanovesicle samples. The metabolites present in the BMNV samples were analyzed via LC–QTOF–MS in both positive and negative ionization modes. The resulting chromatograms are presented in Supplementary Figure 2, while the full metabolite annotation list is included in Supplementary File 1. A total of 163 metabolites were detected. An extensive diversity of metabolites was observed, with nearly one hundred metabolite classes detected in BMNVs. (Figure 4a and Supplementary File 1).

**Figure 4.**
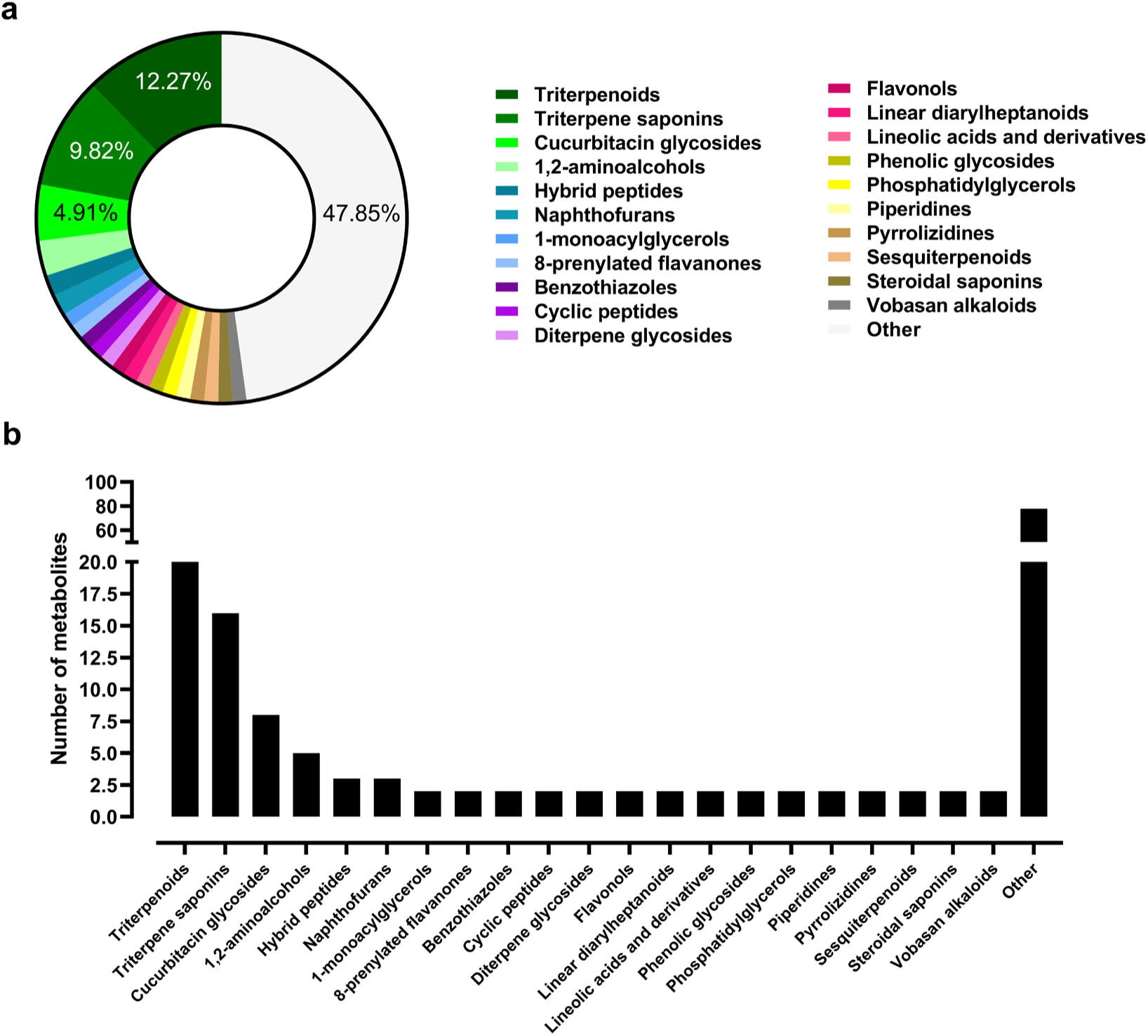
Various classes of the identified metabolites in BMNVs. **(a)** Categories of metabolite classes identified in BMNV samples by LC–QTOF–MS analysis. Each color denotes a distinct metabolite class, and the corresponding area represents the relative percentage of each class. **(b)** A bar graph illustrating the number of identified compounds across different metabolite classes.

Interestingly, among the identified metabolites, triterpenoids and triterpene saponins constituted the predominant chemical classes, reflecting the intrinsic phytochemical profile of the plant, which is known to be enriched in these compounds (Figure 4b). However, bacoside, bacopaside and bacopasaponin were not found. The identified triterpenoids included scabioside C, ursolic acid, echinocystic acid, ganoderic acid LM2, wilforlide A, 16β-hydroxytrametenolic acid, glycyrrhetinic acid, saikogenin D, kalopanaxsaponin H, ganoderic acid G, ginsenoside Rg6, and cauloside C. In addition, several triterpene saponins, including asperosaponin VI, notoginsenoside R1, diacetylpyxinol, ecliptasaponin D, saikosaponin B3, madecassoside, and ginsenoside compound K, were identified.

### 3.3 Superoxide dismutase–related proteins were identified as constituents of the BMNV cargo

Protein identification was conducted to profile the list of proteins in BMNVs. The *B. monnieri* proteome has not yet been fully studied. Therefore, a short list of proteins was identified, and 50.94% of the identified proteins were labeled uncharacterized (Figure 5a). A total of 5 protein groups were identified when searching against the *Bacopa* genus database. Owing to the incomplete protein database of Bacopa species, the Plantaginaceae database was used, and 27 protein groups were detected (Supplementary File 2).

**Figure 5.**
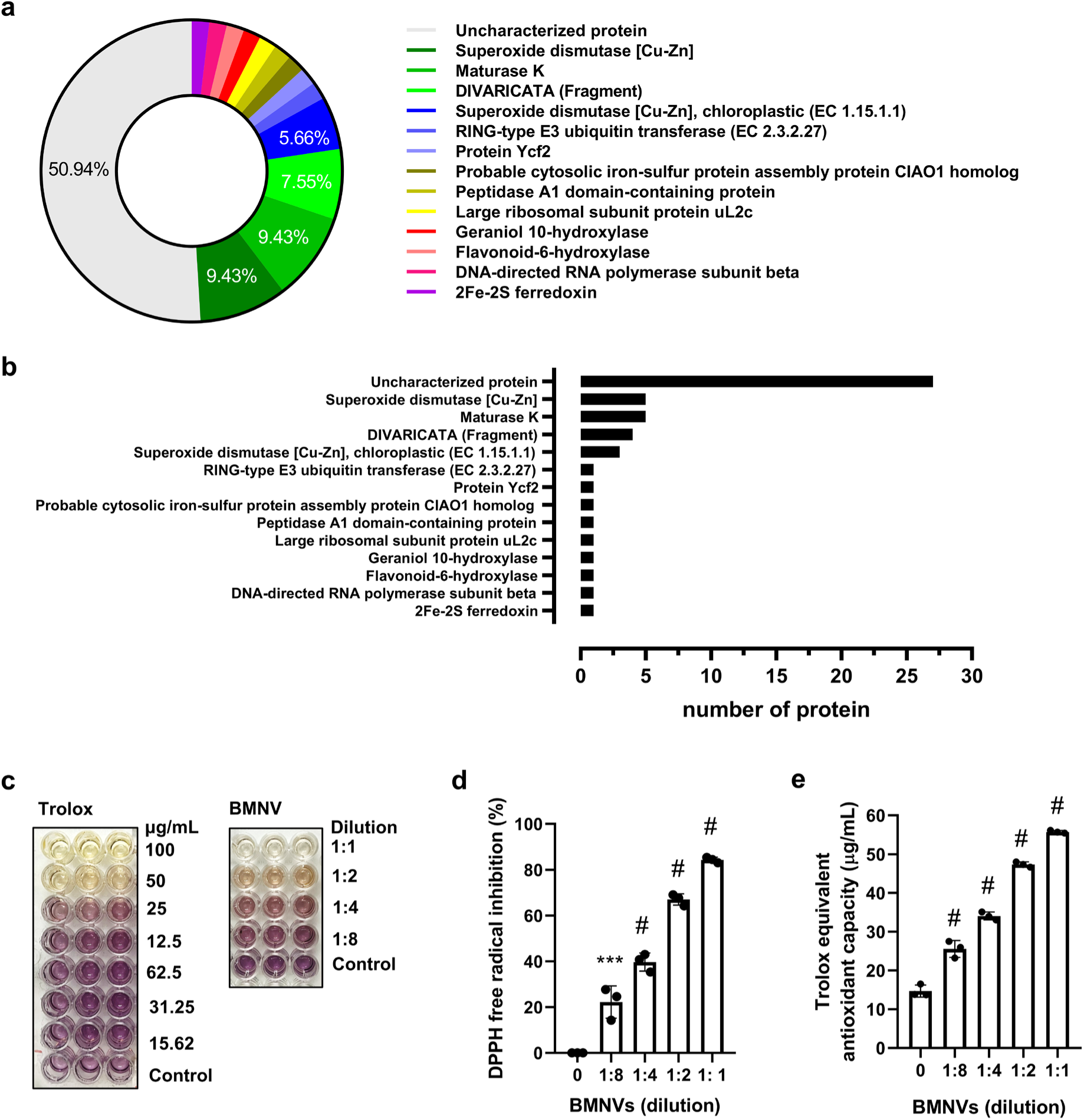
List of the proteins identified in BMNVs and their free radical scavenging potential. **(a)** Distribution of protein groups identified in BMNV samples via LC‒MS analysis. Each color denotes a distinct protein class, and the corresponding area represents the relative proportion of each class. **(b)** Bar graph showing the number of proteins identified within each protein group. **(c)** Representative images of the DPPH assay results illustrating intensity discoloration, demonstrating the free radical scavenging activity of Trolox (standard antioxidant) and BMNVs at different concentrations (μg/mL) and dilutions (MBNV and methanol ratio), respectively. **(d)** Quantitative analysis of the DPPH free radical scavenging activity (%) of BMNVs at the indicated dilutions. **(e)** Trolox equivalent antioxidant capacity (TEAC) of BMNVs expressed as μg/mL Trolox equivalents. The data are presented as the means ± SDs, and individual dots represent independent measurements. Statistical significance among experimental groups was determined using one-way ANOVA followed by Tukey’s multiple comparisons test. *** p<0.001, # P<0.0001 (vs group 0).

Unexpectedly, although only a limited number of annotated proteins were detected in the BMNV samples, superoxide dismutase (SOD)-related proteins constituted the predominant protein groups. Specifically, SOD accounted for 9.43% of the identified proteins, whereas chloroplast-localized SOD represented an additional 5.66%, ranking these proteins among the top two most abundant protein families detected, with identification scores exceeding 10.0. The data also indicate that a total of eight SOD-related protein entries (three chloroplastic and five cytosolic) were detected in the BMNV samples (Figure 5b). Other classes of annotated proteins include maturase K, DIVARICATA (Fragment), RING-type E3 ubiquitin transferase (EC 2.3.2.27), protein Ycf2, probable cytosolic iron‒sulfur protein assembly protein CIAO1 homolog, peptidase A1 domain-containing protein, large ribosomal subunit protein uL2c, geraniol 10-hydroxylase, flavonoid-6-hydroxylase, DNA-directed RNA polymerase subunit beta, and 2Fe‒2S ferredoxin.

Despite the overall low number of annotated proteins detected in BMNVs, the relatively high representation of SOD isoforms underscores the enrichment of antioxidant-related proteins within these plant-derived nanovesicles. Therefore, the antioxidant properties of the isolated BMNVs were further evaluated by assessing free radical scavenging activity via the DPPH assay. The results demonstrated that the BMNVs effectively neutralized DPPH free radicals, as evidenced by a visible color change from dark violet to yellow across all the tested concentrations (Figure 5c). The percentage of free radical inhibition reached a maximum of over 80%, suggesting the potent antioxidant activity of the BMNVs (Figure 5d). BMNVs clearly exhibited a dose-dependent increase in antioxidant capacity, as quantified by the Trolox equivalent antioxidant capacity (TEAC) (Figure 5e). At higher concentrations, BMNVs presented a marked increase in antioxidant capacity, reaching approximately 50 µg/mL Trolox equivalents at a 1:2 dilution.

### 3.4 BMNV treatment suppressed neuroblastoma cell growth and elicited morphological changes that differed from the cytotoxic effects induced by doxorubicin

To evaluate the potential bioactivities of the isolated BMNVs, their growth-suppressive effects were examined in the human neuroblastoma SH-SY5Y cell line and the non-tumorigenic human kidney HEK293 cell line. Cell viability was assessed via the MTT assay. Doxorubicin, an FDA-approved chemotherapeutic agent for neuroblastoma, was used as a positive control [30].

Doxorubicin (5 μM) suppressed the growth of both SH-SY5Y and HEK293 cells. In the SH-SY5Y cells, sublethal doses of BMNVs were detected at concentrations lower than 5x 10^7^ particles/mL (Figure 6a). The highest concentration (5x 10^10^ particles/mL) reduced viability to approximately 42%, achieving a level of inhibition comparable to that of the doxorubicin-treated positive control. While this reduction is statistically significant, the retention of nearly half the cell population suggests a moderate cytotoxic or cytostatic effect rather than acute, nonspecific toxicity. This observation implies that BMNVs do not induce immediate cell death. Crucially, BMNVs demonstrated a favorable safety profile, with no significant toxicity observed in HEK293 cells at doses (5 x10^8^ and 5 x 10^9^ particle/mL) that were cytotoxic to neuroblastoma cells (Figure 6a), suggesting their potential as selective nanotherapeutic agents that target rapidly dividing malignant cells.

**Figure 6.**
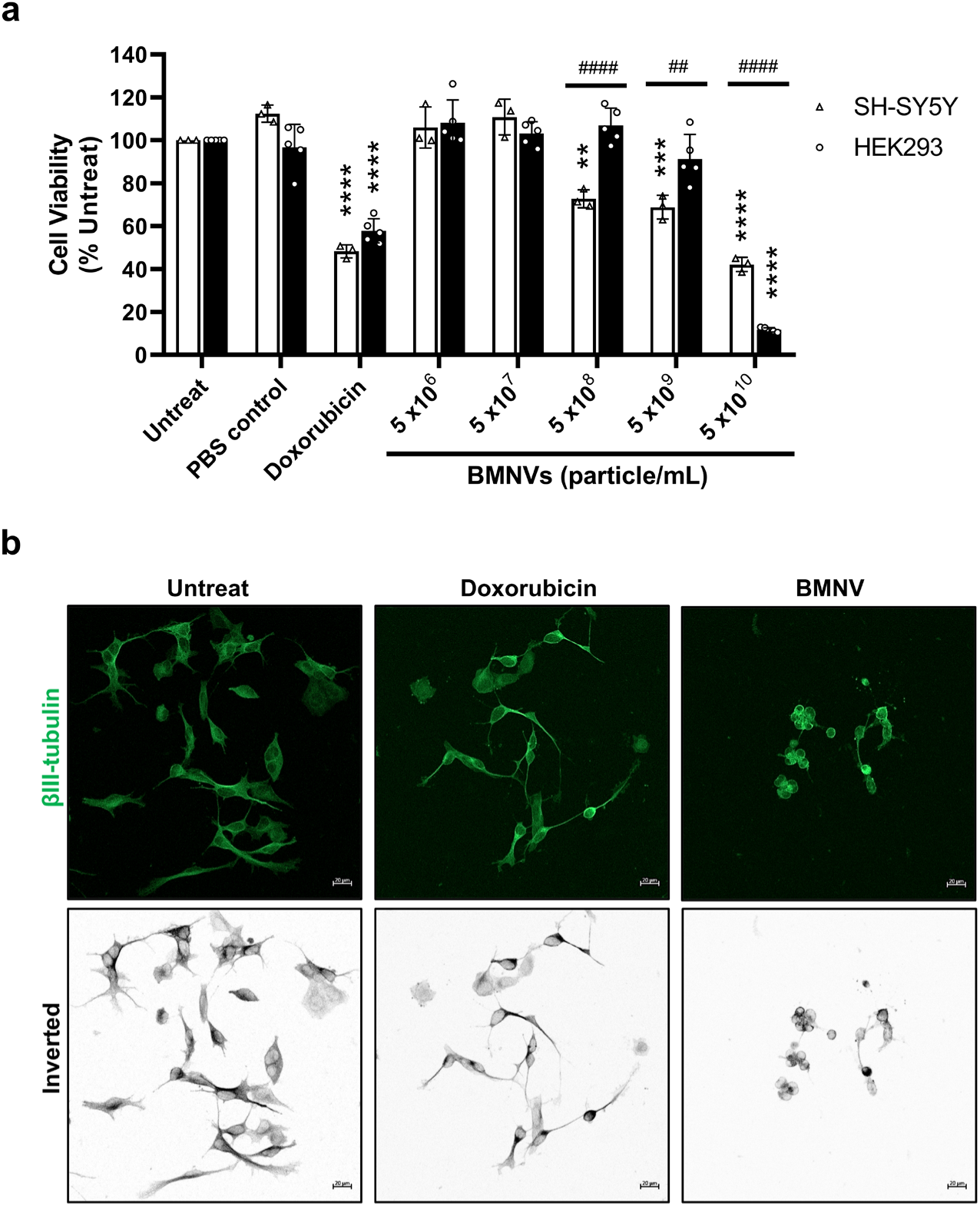
BMNVs suppress neuroblastoma cell growth. **(a)** Quantitative analysis of cell viability was assessed via the MTT assay in human SH-SY5Y and HEK293 cells following exposure to doxorubicin (5 µM) and increasing concentrations of BMNVs (particles/mL) for 24 h, relative to untreated controls. Data are expressed as percentages normalized to the untreated control and presented as the means ± SDs from different biological replicates (n = 3–6). Statistical significance among experimental groups was determined using two-way ANOVA. ** p<0.01, ***p < 0.001, ****p < 0.0001 (vs. untreat). ## p<0.01, #### p<0.0001 (SH-SY5Y vs HEK293). **(b)** Representative immunofluorescence images of SH-SY5Y cells stained for βIII-tubulin (green) under untreated, doxorubicin (5 μM), and BMNV (5x 10^8^ particles/mL)-treated conditions. The corresponding inverted grayscale images are shown to enhance the visualization of the cellular structure. Scale bars = 20 µm.

Notably, the decrease in cell viability appeared to be linked to growth arrest rather than overt cytotoxicity. Figure 6b highlights clear differences in morphological outcomes between doxorubicin and BMNV treatments. While doxorubicin induced an elongated, spindle-like morphology with filamentous protrusions indicative of stress-related cytoskeletal alterations, BMNV treatment (5 x 10^8^ particle/mL) resulted in a spherical and shrunken cellular appearance. The divergence in morphological phenotypes supports the notion that BMNVs elicit cellular responses through mechanisms distinct from those of doxorubicin.

### 3.5 BMNV-induced cell arrest at non-G0 phases underlies the mechanism of growth suppression in neuroblastoma SH-SY5Y cells

Doxorubicin is known to activate the DNA damage response, leading to cell cycle arrest, specifically at the G0 phase, in astrocytes and microglia [31,32]. To determine whether doxorubicin induces similar G0 arrest in SH-SY5Y neuroblastoma cells, we performed immunocytochemical staining for Ki-67, a biomarker expressed during all active phases of the cell cycle (G1, S, G2, and M) but absent in G0.

Following doxorubicin treatment, Ki-67 immunoreactivity was abolished, confirming G0 phase arrest. In contrast, the BMNV-treated cells remained Ki-67 positive (Figure 7), indicating that these cells resided within the non-G0 phase. These data indicate that BMNV treatment of SH-SY5Y cells resulted in the accumulation of cells in the S or G2/M phases of the cell cycle. Collectively, these findings further suggest that the anticancer activity of BMNVs is mediated through a mechanism distinct from that of conventional chemotherapeutic agents.

**Figure 7.**
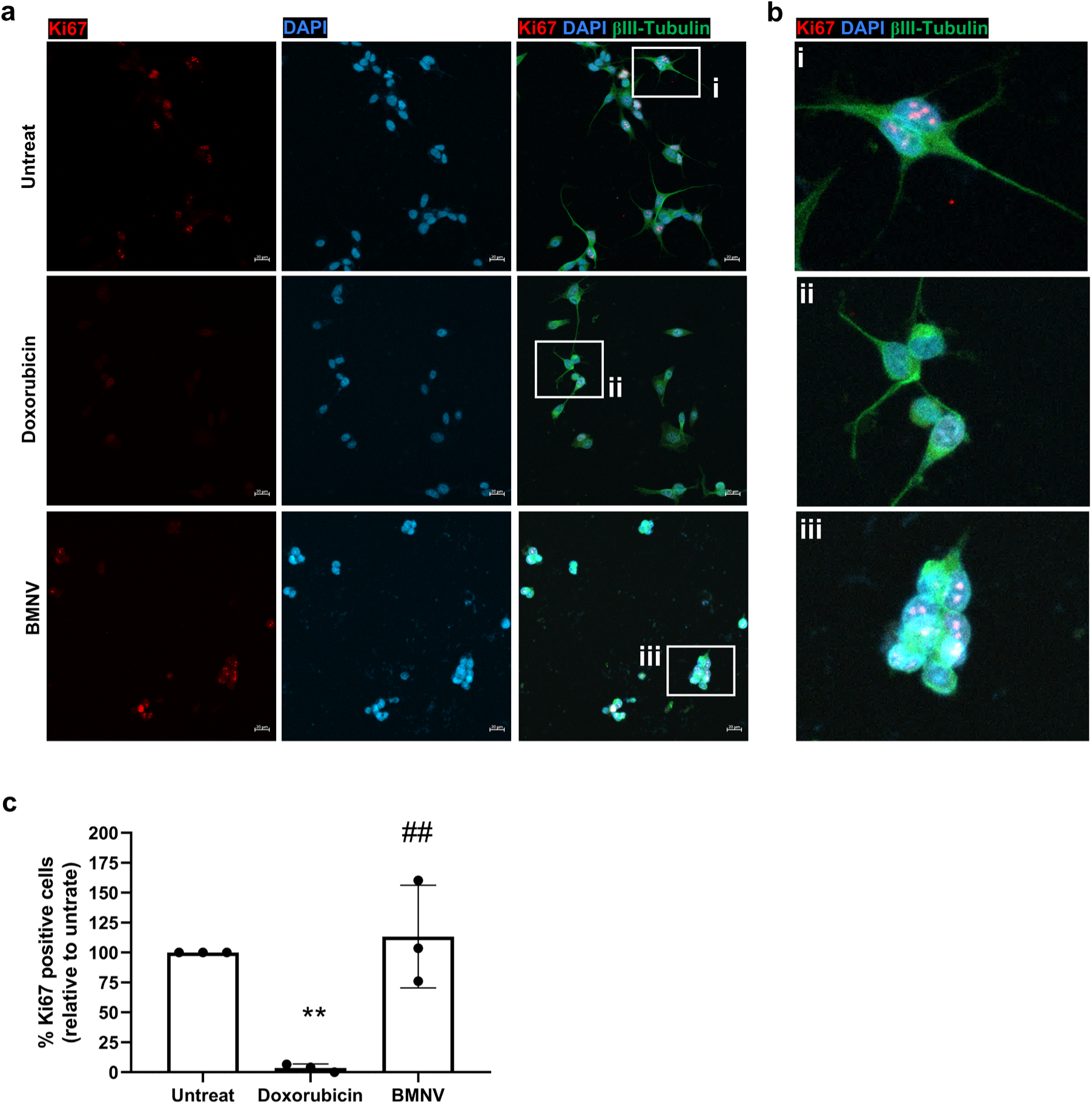
The effect of BMNVs on Ki67 expression in SH-SY5Y cells. **(a)** Representative immunofluorescence images of SH-SY5Y cells under untreated, doxorubicin-treated (5 µM), and BMNV (5x 10^8^ particles/mL)-treated conditions, stained for Ki67 (red), βIII-tubulin (green), and nuclei (DAPI, blue). The scale bar is 20 μm. **(b)** Higher-magnification images corresponding to the boxed regions in panels (a) (i–iii), highlighting Ki67-positive nuclei in untreated cells, marked suppression of Ki67 expression following doxorubicin treatment, and the presence of Ki67-positive cells with BMNV exposure. **(c)** Quantitative analysis of Ki67-positive cells expressed as a percentage of total DAPI-stained nuclei. The data are presented as the means ± SDs from three biological replicates. Statistical significance is indicated as **p < 0.01 (vs the untreated group) and ## p < 0.01 (vs the doxorubicin-treated group).

### 3.6 Cellular internalization of BMNVs in neuroblastoma SH-SY5Y cells

To investigate the cross-kingdom interaction between plant-derived nanovesicles and human neuroblastoma cells, a cellular uptake assay was conducted. BMNVs were labeled with the lipophilic dye PKH26 and incubated with SH-SY5Y cells to track vesicle internalization. Two-dimensional fluorescence imaging revealed significant accumulation of punctate fluorescent signals within the SH-SY5Y cytoplasmic compartment (Figure 8a-c). Furthermore, three-dimensional reconstruction confirmed the predominantly perinuclear localization of BMNVs within these cells (Figure 8d and Supplementary File 3). These results demonstrate that BMNVs are efficiently internalized by SH-SY5Y cells, providing a mechanistic basis for the observed intracellular effects. The efficient internalization of BMNVs highlights that they could also be developed as promising anticancer drug delivery platforms targeting SH-SY5Y neuroblastoma.

**Figure 8.**
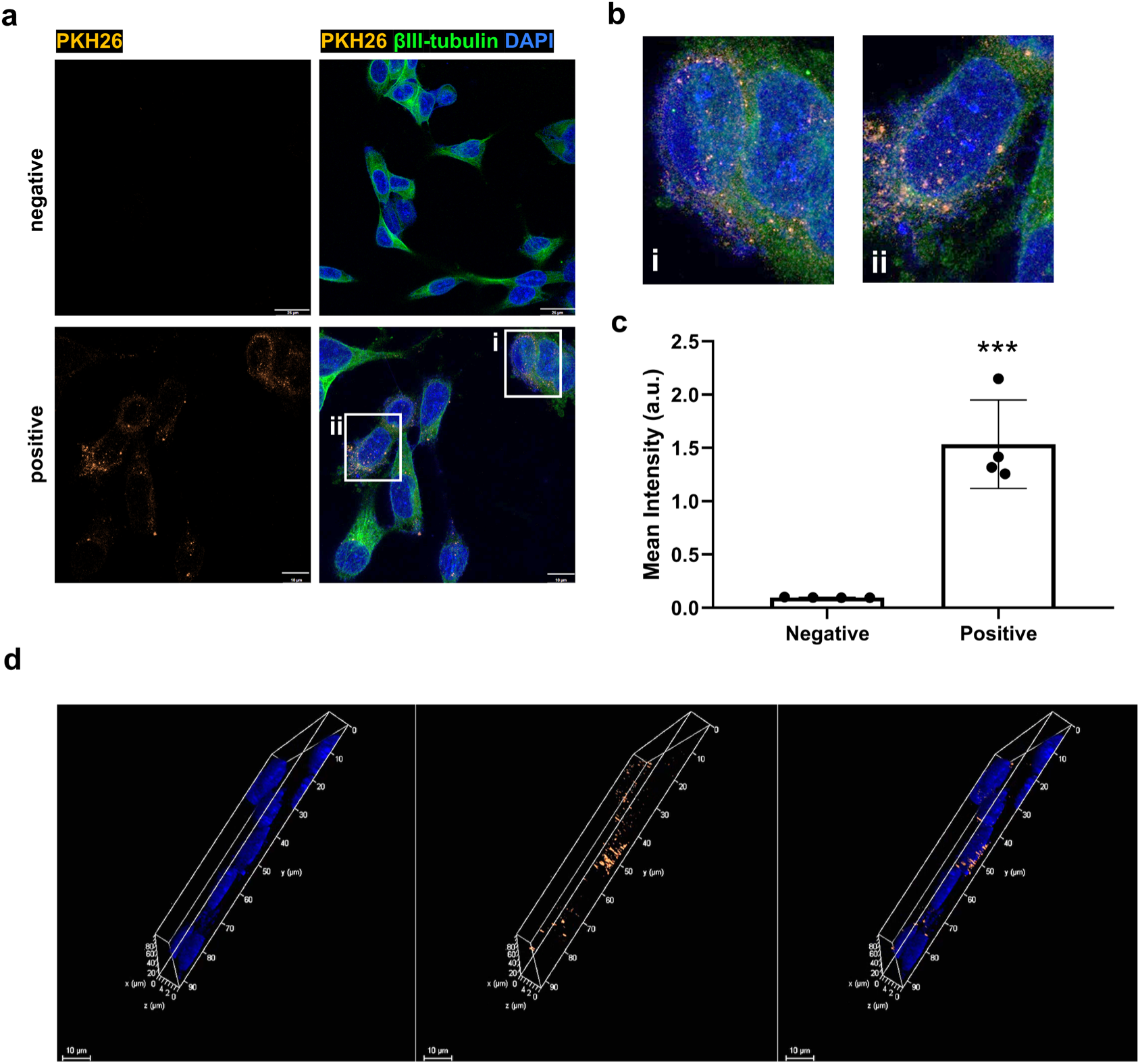
The internalization of BMNVs in neuroblastoma SH-SY5Y cells. **(a)** PKH26 labeling combined with immunocytochemistry for βIII-tubulin (a neuronal lineage marker) and nuclear staining (DAPI) demonstrated the internalization of BMNVs in SH-SY5Y cells. Scale bar: 10 μm. **(b)** Enlarged 2D image from the square area highlighted in (a) showing PKH26-labeled nanovessicles localized within neuroblastoma cells. **(c)** Quantification of the PKH26 fluorescence intensity compared between PKH26-negative and PKH26-positive conditions. The data are presented as the means ± SDs. Statistical significance is indicated as ***p < 0.001. **(d)** 3D reconstruction confirming the intracellular internalization and perinuclear distribution of the BMNVs in neuroblastoma cells. Scale bar: 10 μm. The 3D video files are provided in Supplementary File 3.

## 4. Discussion

The emergence of plant-derived nanovesicles (PDNVs) as a therapeutic modality offers a unique intersection between traditional ethnopharmacology and modern nanomedicine. In this study, we successfully isolated and characterized nanovesicles from *Bacopa monnieri* (BMNVs), providing a comprehensive analysis of their biophysical properties, molecular cargo, and anticancer potential against human neuroblastoma cells.

The *B. monnieri* plant, commonly known as Brahmi, is a medicinal plant in Ayurvedic medicine that is traditionally used for its neuroprotective, memory-enhancing, and antioxidant properties[15,22]. Its pharmacological activity is attributed mainly to a group of dammarane-type triterpenoid saponins largely bacoside A (mixture), bacopasides, and related triterpenoid saponins, in addition to supportive phenolics and terpenoids[33–35]. Despite their therapeutic potential, bacosides (e.g., Bacoside A) are characterized by poor water solubility, a bitter taste, and low oral bioavailability, which necessitates high dosages and often limits patient compliance[18]. The utilization of BMNVs offers a strategic approach to overcome these pharmacokinetic limitations. As biogenic carriers, PDNVs are known to increase the stability and solubility of hydrophobic phytochemicals, facilitating their transport across biological membranes, including the blood‒brain barrier (BBB), while maintaining low immunogenicity[36–39]. In this study, we successfully isolated BMNVs for the first time, demonstrating that these vesicles are naturally enriched with active phytochemical agents. Our findings establish a novel biotechnological platform through the isolation of BMNVs, effectively transforming *B. monnieri* from a crude herbal extract into a modern nanomedicine. This study highlights the potential for these biogenic vesicles to be explored for several biomedical applications, including the management of diverse neurological disorders.

Ultracentrifugation is still the core method for isolating PDNVs, but coupling ultracentrifugation with additional purification steps markedly improves their purity, reproducibility, and suitability for downstream mechanistic studies. Most PDNV protocols start from plant juice and use differential ultracentrifugation (dUC), which is the benchmark approach because it is scalable, relatively inexpensive per sample, and works for many matrices [40–43]. However, dUC alone often co-sediments proteins, nucleic acids, and matrix components, and high g-forces can induce vesicle aggregation or damage. Currently, extracellular vesicle isolation is performed with combined isolation methods, as combined methods are superior to single EV isolation methods for improving vesicle purity[44,45]. We combined dUC and ultrafiltration (UF) using a Macrosep ultracentrifuge device to concentrate and yield higher-purity BMNVs. UF is pressure-driven, fast, and easy to scale, has less operator dependence than UC does and is more compatible with GMP-like workflows[46–48]. Additionally, UF can wash out small soluble proteins (<100 kDa) while retaining the nanovesicles in the retentate.

Characterization methods for PDNVs differ from those used for mammalian-derived EVs, as the biochemical profiles of specific plant-derived markers are not yet well standardized. Unlike PDNVs, which currently lack universal standardization, mammalian EV markers (e.g., CD63, CD81, and TSG101) are well established by the International Society for Extracellular Vesicles (ISEV)[29,49,50]. No single protein is a universal EV marker. Therefore, multiple markers and methods should always be combined[29]. In this study, the biophysical properties of BMNVs were rigorously established via NTA and Cryo-EM. These techniques confirmed a standardized particle size distribution (∼112 nm) and the preservation of a distinct lipid bilayer morphology, which are critical parameters for ensuring nanovesicle integrity. However, a current limitation in the field remains the standardization of biochemical markers for PDNVs that can serve as definitive standards across different botanical sources.

PDNVs carry proteins, lipids, metabolites and RNAs that can affect mammalian cells[45]. Mass spectrometry (MS) is the central tool for defining the protein and metabolite compositions of PDNVs in an unbiased way [45,51–53]. The enrichment of triterpenoids and triterpene saponins within BMNVs can be explained by several complementary biological and physicochemical factors intrinsic to both the plant and the biogenesis of PDNVs. *B. monnieri* is well recognized for its distinctive secondary metabolite profile, in which triterpenoids and triterpene saponins represent the dominant and most abundant classes of bioactive compounds[12,33,34]. These metabolites are synthesized through the mevalonate pathway and are constitutively produced at relatively high levels in aerial tissues, particularly in leaves and stems, which were the source material used for nanovesicle isolation in this study. Consequently, nanovesicles originating from these tissues are inherently exposed to, and likely to incorporate, these metabolites during vesicle formation. From a mechanistic perspective, PDNVs are formed through endosomal and plasma membrane–associated trafficking pathways that selectively package cellular components, including lipids and secondary metabolites, into lipid bilayer structures. Triterpenoids and triterpene saponins possess amphiphilic properties characterized by hydrophobic aglycone cores and, in the case of saponins, hydrophilic sugar moieties[54]. This structural feature strongly favors their association with lipid membranes, facilitating their preferential partitioning into nanovesicular bilayers during vesicle biogenesis.

Although bacoside A was not detected in the metabolite profile of BMNVs, several of the identified metabolites are well documented for their anticancer activities (Supplementary Table 1). Notable examples include flavonols (such as kaempferol)[55], saikosaponins [56–58], madecassoside [59,60], ginsenoside compound K [61,62], ursolic acid [63,64], and echinocystic acid [65]. In addition to their anticancer properties, a number of these metabolites have also been reported to exert protective effects across diverse disease models. Madecassoside, one of the four major triterpenoids in *Centella asiatica*, has been extensively studied for its protective effects [66–69]. Several metabolites identified in BMNVs have been shown to suppress inflammation, including wilforlide A [70], ginsenoside Rg6 [71], kaempferol [72], kalopanax saponin H [73], glycyrrhetinic acid [74], and ganoderic acid G [75]. Collectively, these metabolites exert anti-inflammatory effects primarily by downregulating proinflammatory cytokines and signaling pathways such as NF-κB, suggesting that BMNVs may warrant future investigations as potential natural anti-inflammatory agents.

Protein profiling of BMNVs was also performed, yielding a relatively short list of confidently identified proteins. A substantial proportion of these proteins were classified as uncharacterized (50.94%), which can be attributed to incomplete genomic sequencing and the limited proteomic annotation available for *B. monnieri*, thereby restricting the representation of its protein sequences in commonly used reference databases. This constraint is not unique to the present study and reflects a broader challenge in plant-derived nanovesicle research. For example, a quantitative LC–MS/MS analysis of nanovesicles isolated from *Malus domestica* identified 187 proteins, of which approximately 37.5% were similarly annotated as uncharacterized[76]. These observations highlight the current limitations of plant proteomic databases and underscore the need for more comprehensive genomic and proteomic resources to enable deeper characterization of the PDNV protein cargo. Nevertheless, the present study provides compelling evidence for the presence of superoxide dismutase (SOD) within BMNVs, identified with a high confidence score (78.24%), underscoring the intrinsic antioxidant signature of their botanical origin. Although SOD has not been commonly reported as a molecular cargo in previously characterized PDNV proteomic profiles, its presence has been documented in extracellular vesicles isolated from potato (*Solanum tuberosum cv. Laura*) roots [77]. The identification of SOD in BMNVs is consistent with the pronounced free radical–scavenging activity observed in our antioxidant assay. These findings suggest that the isolated nanovesicles may be further developed as cytoprotective agents in disease models driven by oxidative stress, including neurodegenerative disorders. In addition, their antioxidant capacity makes BMNVs attractive candidates for translational biocosmetic applications[78]. Additionally, superoxide dismutase can play context-dependent roles in cancer and has, in some settings, been implicated in tumor-promoting redox signaling[79,80]. Therefore, the presence of SOD within BMNVs may contribute to altered redox homeostasis in neuroblastoma cells and could partly underlie the anti-neuroblastoma effects observed in this study, although further mechanistic validation is needed.

The potential bioactivities of BMNVs were evaluated in neuroblastoma SH-SY5Y cells, where they suppressed cell growth and induced marked alterations in cellular morphology, whereas even at toxic doses, BMNVs did not significantly reduce cell viability in the non-tumorigenic human kidney HEK293 cell line. This finding is consistent with multiple studies on plant-derived nanovesicles, which have reported stronger cytotoxic effects in cancer cells than in normal fibroblasts or other nonmalignant cell types [81–83]. The observation that BMNVs induce cell cycle arrest in non-G0 phases suggests a mechanism of action (MOA) distinct from that of the conventional chemotherapeutic drug doxorubicin, which typically triggers robust exit from the cell cycle, leading to G0-phase arrest or senescence [29–31]. This finding is consistent with reports on other plant-derived nanovesicles. For example, ginger exosome-like nanovesicles have been shown to suppress the growth of human breast adenocarcinoma cells (MDA-MB-231) by inducing G2/M phase arrest through the downregulation of cyclin B and Cdc2 [8]. Grapefruit-derived nanovesicles inhibited the proliferation of A375 melanoma cells by arresting the cells at the G2/M checkpoint, resulting in the downregulation of cyclins B1/B2 and the upregulation of p21[84]. Therefore, these intrinsic anticancer activities occur via cell cycle arrest at non-G0 phases, depending on the plant source and tumor type. While the present findings indicate that BMNVs interfere with cell proliferation, further studies using both *in vitro* and *in vivo* models, incorporating comprehensive molecular analyses of key cell cycle regulators, are needed to fully elucidate the signaling pathways modulated by BMNVs.

The cellular uptake of PDNVs by mammalian recipient cells has been extensively investigated as a key line of evidence supporting cross-kingdom interactions. The efficient uptake of BMNVs by neuroblastoma SH-SY5Y cells demonstrated in the present study is consistent with a substantial body of published evidence showing that PDNVs can be readily internalized and exert intracellular biological effects in mammalian systems[82,85]. These findings further support the intrinsic biocompatibility of PDNVs and underscore their suitability for development as natural delivery platforms for bioactive compounds. Moreover, three-dimensional confocal imaging revealed a distinct perinuclear localization of BMNVs within SH-SY5Y cells. This distribution strongly suggests the occurrence of intracellular trafficking rather than nonspecific adsorption to the plasma membrane. Notably, a comparable perinuclear localization pattern was observed in our previous studies examining the internalization of human stem cell–derived exosomes in central nervous system mouse neuronal cells, further reinforcing the notion that vesicle uptake and intracellular trafficking mechanisms are conserved across different vesicle origins and recipient cell types [86].

Our work provides the first evidence for the isolation of *B. monnieri*-derived nanovesicles enriched with bioactive cargo. The attenuation of neuroblastoma growth underscores the bioactive potential of the nanovesicles and supports their prospective development as plant-derived nanotherapeutic agents for cancer therapy. A diagram summarizing the key findings of this work is shown in Figure 9.

**Figure 9.**
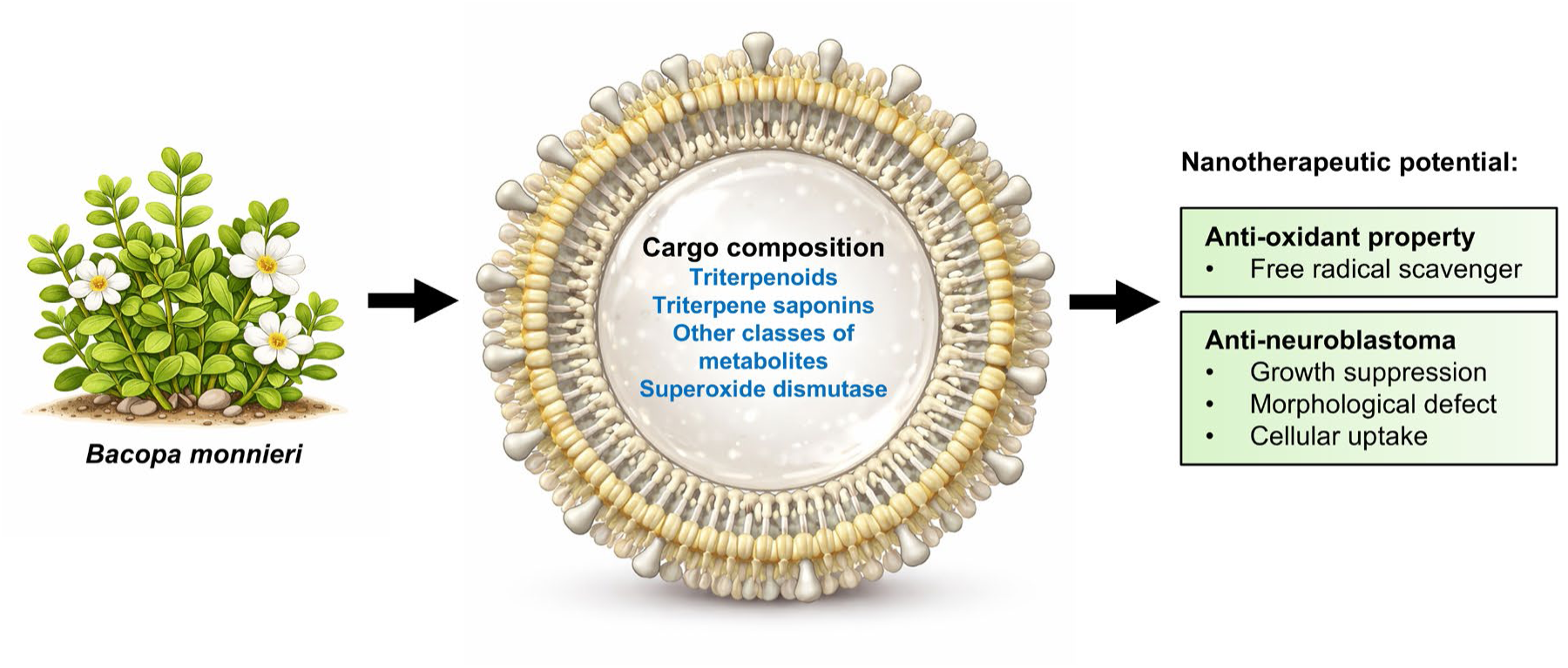
Schematic summary of the key findings in this study.

## Declaration of generative AI and AI-assisted technologies in the writing process

This manuscript was prepared with the assistance of generative AI and AI-assisted technologies to improve its clarity, grammar, and overall readability. After using these tools, the authors reviewed and edited the content. Curie was employed for language editing and text generation.

## Author contributions

**Ekkaphot Khongkla:** Conceptualization, methodology, validation, formal analysis, investigation, writing - original draft, visualization, and funding acquisition. **Panitch Boonsnongcheep:** Investigation, Formal analysis, Resources, Supervision, Writing - Review & Editing. **Pipob Suwanchaikasem:** Investigation, Formal analysis. **Kornkanok Promthep**: Investigation. **Monruedee Srisaisup and Theptharin Charuraksa**: Investigation, Formal analysis. **Pannaphan Makarathut:** Investigation, Formal analysis. **Banthit Chetsawang:** Resources, Supervision, Writing - Review & Editing.

## Acknowledgment

The authors thank the Mahidol University Frontier Research Facility (MU-FRF) for their technical assistance with Cryo-EM service.

## Data availability

The data will be made available upon proper request to the corresponding author.

