## Supplementary Information for "*Bacopa monnieri (L.)* Wettst-derived nanovesicles are enriched with bioactive cargo and exhibit anti-neuroblastoma activity"

### Isolation and characterization of *Bacopa monnieri* (L.) Wettst-derived nanovesicles with anti-neuroblastoma potential

#### Supplementary Files

**Supplementary File 1.** The full list of identified metabolites in BMNVs.

**Supplementary File 2.** The full list of identified proteins in BMNVs.

**Supplementary File 3.** The video visualizing perinuclear distribution of BMNVs in neuroblastoma SH-SY5Y cells.

**Supplementary Figures**

**Supplementary Figure 1.** NTA-based stability analysis of BMNVs. Comparison of **(a)** size distribution and **(b)** particle concentration of BMNVs prior to and following six weeks of storage at -30°C. Data are presented as the mean ± SDs. Statistical significance was determined using a Student's t-test showing not significant.

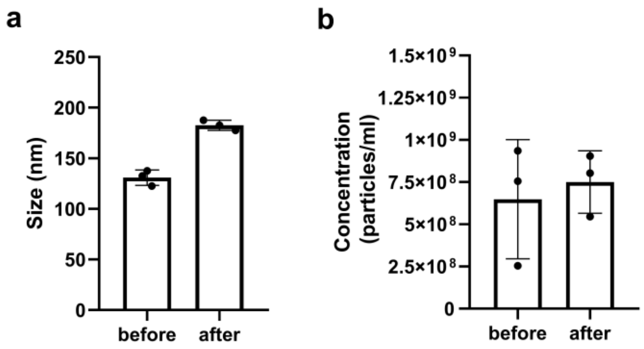

**Supplementary Figure 2.** Total ion chromatograms derived from metabolite identification analyses in **(a)** positive and **(b)** negative ionization modes. The signal intensity (a.u.) of the blank control (black trace) and MBNVs (red trace) is displayed across the acquisition time (min).

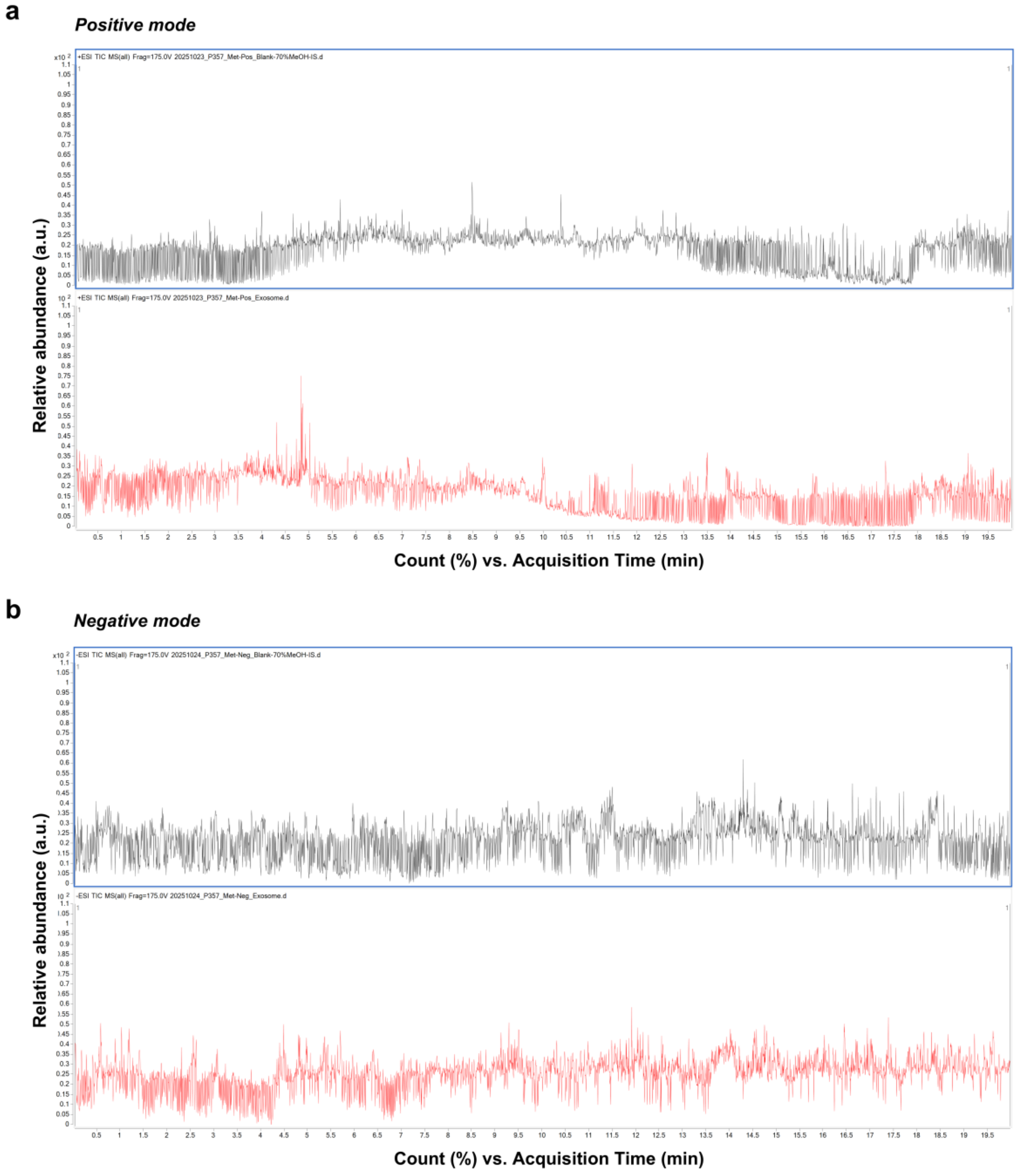

**Supplementary Figure 3.** Total ion chromatograms derived from peptide identification analyses. The signal intensity (a.u.) of the blank control (black trace) and MBNVs (red trace) is displayed across the acquisition time (min).

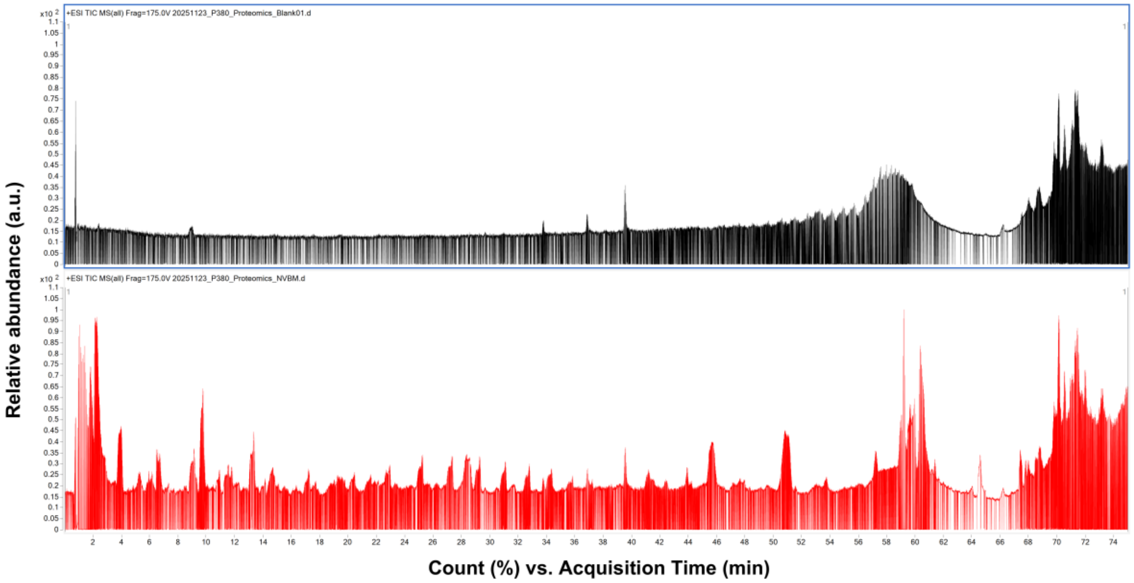

#### Supplementary Tables

#### **Supplementary Table 1.** List of metabolites identified in *B. monnieri* -derived nanovesicles

| No. | Metabolite name | Ontology | Average RT (min) | Average Mz |
|---|---|---|---|---|
| 1 | Lauryldiethanolamine | 1,2-aminoalcohols | 9.791 | 274.27405 |
| 2 | D-Sphingosine | 1,2-aminoalcohols | 11.493 | 300.29062 |
| 3 | 2,2'-(Tetradecylimino)diethanol | 1,2-aminoalcohols | 11.237 | 302.30566 |
| 4 | Dehydrophytosphingosine (not validated, isomer of 1677) | 1,3-aminoalcohols | 10.867 | 316.28568 |
| 5 | D-ribo-Phytosphingosine | 1,3-aminoalcohols | 9.917 | 318.30093 |
| 6 | Gliquidone | 1,3-isoquinolinediones | 4.861 | 550.20087 |
| 7 | LPC 18:2 | 1-acyl-sn-glycero-3-phosphocholines | 12.218 | 520.34058 |
| 8 | 2,3-dihydroxypropyl stearate | 1-monoacylglycerols | 15.118 | 381.29764 |
| 9 | 2,3-dihydroxypropyl palmitate | 1-monoacylglycerols | 14.594 | 353.26682 |
| 10 | MMV676382 | 3-alkylindoles | 4.861 | 378.20325 |
| 11 | TOBRAMYCIN | 4,6-disubstituted 2-deoxystreptamines | 14.120 | 485.28870 |
| 12 | Eupatilin | 6-O-methylated flavonoids | 8.226 | 345.08737 |
| 13 | 6,8-Diprenylnaringenin | 8-prenylated flavanones | 5.544 | 407.18460 |
| 14 | NCGC00385984-01!2-(3,5-dihydroxyphenyl)-5,7-dihydroxy-6,8-bis(3-methylbut-2-enyl)-2,3-dihydrochromen-4-one | 8-prenylated flavanones | 4.058 | 423.17914 |
| 15 | Lasiocarpine N-oxide | Alkaloids and derivatives | 6.456 | 428.23117 |
| 16 | Valganciclovir | Alpha amino acid esters | 14.875 | 353.16394 |
| 17 | Dimethachlor OXA | Alpha amino acids and derivatives | 7.028 | 250.10823 |
| 18 | Putative dehydroxy-amphotericin B | Aminoglycosides | 4.156 | 908.48810 |
| 19 | Pyrenocine B putative | Aryl alkyl ketones | 13.177 | 227.08609 |
| 20 | NCGC00385365-01_C11H16O3_2(4H)-Benzofuranone, 5,6,7,7a-tetrahydro-6-hydroxy-4,4,7a-trimethyl-, (6S,7aR)- | Benzofurans | 8.876 | 219.09567 |
| 21 | 2-Mercaptobenzothiazole | Benzothiazoles | 13.497 | 167.99393 |
| 22 | 2-(4-Morpholinyl)benzothiazole | Benzothiazoles | 7.725 | 221.07420 |
| 23 | NCGC00180625-02!3-[5,7-dihydroxy-2-(4-hydroxyphenyl)-4-oxo-2,3-dihydrochromen-3-yl]-5,7-dihydroxy-2-(4-hydroxyphenyl)-2,3-dihydrochromen-4-one | Biflavonoids and polyflavonoids | 4.672 | 587.11823 |
| 24 | "(3S,5R,7R,8R,9S,10S,13S,14S,16R,17R)-17-((2R)-7-acetoxy-6-methylheptan-2-yl)-10,13-dimethylhexadecahydro-1H-cyclopenta[a]phenanthrene-3,7,16-triyl triacetate" | Bile acids, alcohols and derivatives | 10.654 | 605.40662 |
| 25 | methyl 2-(8-methyl-3a,4,5,6-tetrahydro-1H-pyrazino[3,2,1-jk]carbazol-3(2H)-yl)acetate | Carbazoles | 9.870 | 299.18420 |
| 26 | N-lauroylethanolamine | Carboximidic acids | 8.450 | 244.22783 |
| 27 | 4-((9S,10R,17S)-10-((E)-(decylimino)methyl)-3-(((2R,4S,5S,6R)-4,5-dihydroxy-6-methyltetrahydro-2H-pyran-2-yl)oxy)-5,14-dihydroxy-13-methylhexadecahydro-1H-cyclopenta[a]phenanthren-17-yl)furan-2(5H)-one | Cardenolide glycosides and derivatives | 14.497 | 674.46356 |

| | | | | |
|---|---|---|---|---|
| 28 | 4,4'-((9S,9'S,10R,10'R,17S,17'S)-((1E,1'E)-(butane-1,4-diylbis(azanylylidene))bis(methanylylidene))bis(3,5,14-trihydroxy-13-methylhexadecahydro-1H-cyclopenta[a]phenanthrene-17,10-diyl))bis(furan-2(5H)-one) | Cardenolides and derivatives | 8.625 | 883.50775 |
| 29 | quinidine | Cinchona alkaloids | 5.176 | 347.16779 |
| 30 | 1-(3,4-Dimethoxycinnamoyl)piperidine | Cinnamic acids and derivatives | 8.071 | 276.15698 |
| 31 | NCGC00385003-01!4-[3-methyl-5-(5,6,7-trihydroxy-1,2,4a,5-tetramethyl-3,4,6,7,8,8a-hexahydro-2H-naphthalen-1-yl)pentoxy]-4-oxobutanoic acid [IIN-based: Match] | Colensane and clerodane diterpenoids | 14.594 | 425.28650 |
| 32 | [(E,6R)-6-hydroxy-6-[(2S,8S,9R,10R,13R,14S,16R)-16-hydroxy-4,4,9,13,14-pentamethyl-3,11-dioxo-2-[(2S,3R,4S,5S,6R)-3,4,5-trihydroxy-6-(hydroxymethyl)oxan-2-yl]oxy-2,7,8,10,12,15,16,17-octahydro-1H-cyclopenta[a]phenanthren-17-yl]-2-methyl-5-oxohept-3-en-2-yl] acetate | Cucurbitacin glycosides | 8.460 | 743.36322 |
| 33 | (2S,3R,4S,5S)-2-(((2S,3R,4S,5R)-2-(((4R,5aS,7S,9S,11aR,12aS)-3-((2R,5R)-5,6-dihydroxy-6-methylheptan-2-yl)-4,7-dihydroxy-2a,5a,8,8-tetramethylhexadecahydrocyclopenta[a]cyclopropa[e]phenanthren-9-yl)oxy)-4,5-dihydroxytetrahydro-2H-pyran-3-yl)oxy)tetrahydro-2H-pyran-3,4,5-triol | Cucurbitacin glycosides | 9.103 | 779.45984 |
| 34 | Astragaloside II | Cucurbitacin glycosides | 14.832 | 827.49146 |
| 35 | 3,4,5-trihydroxy-6-(hydroxymethyl)oxan-2-yl (2E,6R)-6-[(1R,3R,6S,8R,12S,15R,16R)-13,17-dihydroxy-7,7,12,16-tetramethyl-6-{[(2R)-3,4,5-trihydroxy-6-{[(3,4,5-trihydroxyoxan-2-yl)oxy]methyl}oxan-2-yl]oxy}pentacyclo[9.7.0.0?,?.0?,?.0??,??]octadecan-15-yl]-2-methylhept-2-enoate | Cucurbitacin glycosides | 8.297 | 967.48755 |
| 36 | Cimiracemoside D | Cucurbitacin glycosides | 6.007 | 701.39282 |
| 37 | 3,4,5-trihydroxy-6-(hydroxymethyl)oxan-2-yl (2E,6R)-6-[(1R,3R,6S,8R,12S,15R,16R)-13,17-dihydroxy-7,7,12,16-tetramethyl-6-{[(2R)-3,4,5-trihydroxy-6-{[(3,4,5-trihydroxyoxan-2-yl)oxy]methyl}oxan-2-yl]oxy}pentacyclo[9.7.0.0?,?.0?,?.0??,??]octadecan-15-yl]-2-methylhept-2-enoate | Cucurbitacin glycosides | 9.361 | 943.48804 |
| 38 | NCGC00381046-01_C41H66O14_9,19-Cyclolanost-24-en-26-oic acid, 3-[(2-O-hexopyranosylpentopyranosyl)oxy]-12,15-dihydroxy-, (3beta,8xi,9beta,24E)- | Cucurbitacin glycosides | 10.078 | 781.43817 |
| 39 | 3,4,5-trihydroxy-6-(hydroxymethyl)oxan-2-yl (2E,6R)-6-[(1R,3R,6S,8R,12S,15R,16R)-13,17-dihydroxy-7,7,12,16-tetramethyl-6-{[(2R)-3,4,5-trihydroxy-6-{[(3,4,5-trihydroxyoxan-2-yl)oxy]methyl}oxan-2-yl]oxy}pentacyclo[9.7.0.0?,?.0?,?.0??,??]octadecan-15-yl]-2-methylhept-2-enoate | Cucurbitacin glycosides | 8.293 | 943.48981 |
| 40 | Cucurbitacin E | Cucurbitacins | 10.787 | 579.29352 |
| 41 | (2R,3R,6R)-5-[(1E,3E)-hepta-1,3-dienyl]-2,3-dihydroxy-6-(hydroxymethyl)cyclohexan-1-one | Cyclic alcohols and derivatives | 9.456 | 253.14417 |
| 42 | CYCLOPENTANONE | Cyclic ketones | 3.110 | 102.09157 |
| 43 | Microsporin B | Cyclic peptides | 12.322 | 537.30420 |
| 44 | Microsporin B | Cyclic peptides | 12.507 | 537.30438 |
| 45 | Ketotifen fumarate (Zaditor) | Cycloheptathiophenes | 9.226 | 310.12646 |
| 46 | Simvastatin M+Na | Delta valerolactones | 14.594 | 441.26202 |
| 47 | 4-[4-(1,3-benzodioxol-5-yl)-2,3-dimethylbutyl]-2-methoxyphenol | Dibenzylbutane lignans | 8.900 | 327.18127 |
| 48 | (E)-3-(4-acetyloxy-2,3-dihydroxy-2,5,5,8a-tetramethyl-3,4,4a,6,7,8-hexahydro-1H-naphthalen-1-yl)prop-2-enoic acid | Dicarboxylic acids and derivatives | 8.819 | 377.19434 |

| | | | | |
|---|---|---|---|---|
| 49 | Dehydrotumulosic acid | Dihydroxy bile acids, alcohols and derivatives | 3.284 | 485.35754 |
| 50 | NCGC00381111-01_C23H38N4O6_1-[(2E,4E)-6,7-Dihydroxy-2,4-octadienoyl]prolyl-N-methylvalyl-N~2~-methylalaninamide | Dipeptides | 14.729 | 489.26154 |
| 51 | Zamifenacin | Diphenylmethanes | 4.773 | 416.22815 |
| 52 | 3-[(Z)-5-[6-hydroxy-5,5,8a-trimethyl-2-methylidene-3-[3,4,5-trihydroxy-6-(hydroxymethyl)oxan-2-yl]oxy-3,4,4a,6,7,8-hexahydro-1H-naphthalen-1-yl]-3-methylpent-2-enoxy]-3-oxopropanoic acid | Diterpene glycosides | 7.222 | 569.28918 |
| 53 | MalonylHexsose + Hexsose-deoxyHexose + C20H32 | Diterpene glycosides | 7.222 | 861.41290 |
| 54 | NCGC00385411-01_C24H28O10_Spiro[furan-3(2H),1'(4'H)-naphthalene]-4'a,5'(5'H)-dicarboxylic acid, 6'-(acetyloxy)-5-(3-furanyl)-4,5,6',7',8',8'a-hexahydro-5'-hydroxy-2'-methyl-2-oxo-, dimethyl ester | Diterpene lactones | 6.445 | 521.16687 |
| 55 | Dehydroabietamide | Diterpenoids | 4.672 | 300.19852 |
| 56 | h_163_prostanozol_3'OH | Estrane steroids | 14.497 | 679.41846 |
| 57 | Triethylene glycol bis(2-ethylhexanoate) | Fatty acid esters | 14.594 | 403.30618 |
| 58 | (3R,4S,5S,6R)-2-(6-hydroxy-2,6-dimethylocta-2,7-dienoxy)-6-(hydroxymethyl)oxane-3,4,5-triol | Fatty acyl glycosides of mono- and disaccharides | 5.329 | 355.17328 |
| 59 | 3,4,5-trihydroxy-6-[5-hydroxy-2-(4-hydroxy-3-methoxyphenyl)-3-methoxy-4-oxochromen-7-yl]oxyoxane-2-carboxylic acid | Flavonoid-7-O-glucuronides | 4.861 | 529.09424 |
| 60 | 3,7,4'-Trihydroxyflavone | Flavonols | 7.618 | 269.04449 |
| 61 | Kaempferol | Flavonols | 6.890 | 285.04047 |
| 62 | Octyl gallate | Galloyl esters | 13.177 | 305.13196 |
| 63 | NCGC00385247-01_C21H32O8_(3S,3aS,6E,9S,10E,11aS)-3,6,10-Trimethyl-2-oxo-2,3,3a,4,5,8,9,11a-octahydrocyclodeca[b]furan-9-yl beta-D-glucopyranoside | Germacranolides and derivatives | 5.643 | 430.24420 |
| 64 | Megestrol acetate | Gluco/mineralocorticoids, progestogins and derivatives | 14.973 | 385.23499 |
| 65 | L-Saccharopine | Glutamic acid and derivatives | 9.445 | 277.14197 |
| 66 | ((4R)-4-((3R,5S,7R,9S,10S,12S,13R,14S,17R)-3,7,12-trihydroxy-10,13-dimethylhexadecahydro-1H-cyclopenta[a]phenanthren-17-yl)pentanoyl)valine | Glycinated bile acids and derivatives | 14.136 | 530.34637 |
| 67 | 10-hydroxyusambarine | Harmala alkaloids | 14.124 | 467.28018 |
| 68 | NCGC00386102-01![3-methyl-1-[3-methyl-1-[3-methyl-1-[3-methyl-1-oxo-1-(2,3,4,5-tetrahydroxypentoxy)pentan-2-yl]oxy-1-oxopentan-2-yl]oxy-1-oxopentan-2-yl]oxy-1-oxopentan-2-yl] 2-acetyloxy-3-methylpentanoate | Hexacarboxylic acids and derivatives | 0.768 | 782.44299 |
| 69 | MLS002222329-01!26305-03-3 | Hybrid peptides | 15.681 | 708.46271 |
| 70 | 3-(1H-indol-3-ylmethyl)-6,18-dimethyl-12-(1-phenylethyl)-9,15-di(propan-2-yl)-1,4,7,10,13,16,19-heptazacyclotricosane-2,5,8,11,14,17,20-heptone | Hybrid peptides | 10.078 | 771.40991 |
| 71 | SSR 240612 | Hybrid peptides | 8.484 | 755.34143 |
| 72 | NCGC00179731-03!4-[5-[acetyl(hydroxy)amino]pentylamino]-2-[2-[5-[acetyl(hydroxy)amino]pentylamino]-2-oxoethyl]-2-hydroxy-4-oxobutanoic acid [IIN-based on: CCMSLIB00000847088] | Hydroxy fatty acids | 9.146 | 953.50220 |
| 73 | NCGC00380882-01!5-chloro-2,4-dihydroxy-6-methyl-3-[(E)-3-methyl-5-[(1S,2R,6R)-1,2,6-trimethyl-3-oxocyclohexyl]pent-2-enyl]benzaldehyde [IIN-based: Match] | Hydroxybenzaldehydes | 9.632 | 407.20529 |

| | | | | |
|---|---|---|---|---|
| 74 | NCGC00386096-01!(9E,15E,19E)-3-(2-amino-2-oxoethyl)-11,13-dihydroxy-14,21-dimethoxy-8,10,12-trimethyl-7,21-dioxohenicosa-9,15,19-trienoic acid [IIN-based on: CCMSLIB00000848683] | Hydroxyeicosatrienoic acids | 4.156 | 1101.62512 |
| 75 | coronardine | Ibogan-type alkaloids | 7.671 | 339.17761 |
| 76 | (1r,5R,7S)-2-(1H-indol-4-yl)-5,7-dimethyl-1,3-diazaadamantan-6-one | Indoles | 5.698 | 318.16440 |
| 77 | Genistein | Isoflavones | 7.606 | 271.06033 |
| 78 | (2S,3R)-2-(((S)-7-acetamido-1,2,3-trimethoxy-9-oxo-5,6,7,9-tetrahydrobenzo[a]heptalen-10-yl)amino)-N-(3-methoxypropyl)-3-methyl-N-(((1S,9aR)-octahydro-1H-quinolizin-1-yl)methyl)pentanamide | Isoleucine and derivatives | 14.443 | 743.43457 |
| 79 | NCGC00180578-03!(2S)-2-[2-(1,3-benzodioxol-5-yl)ethyl]-4-methoxy-2,3-dihydropyran-6-one | Kavalactones | 4.861 | 294.13452 |
| 80 | epsilon-Decalactone | Lactones | 14.547 | 171.13805 |
| 81 | NCGC00380242-01_C30H42O9_Card-20(22)-enolide, 1,3-bis(acetyloxy)-14,15-epoxy-7,21-dihydroxy-4,4,8-trimethyl-, (1alpha,3alpha,7alpha,9xi,13alpha,15beta,17alpha)- | Limonoids | 10.808 | 591.27319 |
| 82 | NCGC00347453-02!(E)-1,7-bis(4-hydroxyphenyl)hept-4-en-3-one | Linear diarylheptanoids | 14.606 | 341.14224 |
| 83 | Aceroside VIII | Linear diarylheptanoids | 10.808 | 593.27209 |
| 84 | (10E,15Z)-9,12,13-trihydroxyoctadeca-10,15-dienoic acid | Lineolic acids and derivatives | 8.071 | 351.21460 |
| 85 | FA 18:2+3O | Lineolic acids and derivatives | 8.086 | 327.21753 |
| 86 | 4MeC13SPC | Long-chain fatty acids | 15.374 | 383.18790 |
| 87 | NCGC00381375-01!(6E)-heptadeca-6,16-diene-1,2,4-triol [IIN-based: Match] | Long-chain fatty alcohols | 13.807 | 285.24289 |
| 88 | Alpha-Ergocryptine | Lysergamides | 11.554 | 518.32538 |
| 89 | NCGC00163699-03_C26H34O7_2,4,6,8-Decatetraenedioic acid, mono[5-methoxy-4-[2-methyl-3-(3-methyl-2-buten-1-yl)oxiranyl]-1-oxaspiro[2.5]oct-6-yl] ester, (2E,4E,6E,8E)- | Medium-chain fatty acids | 15.137 | 457.22464 |
| 90 | 5-isopropenyl-2-methyl-2-cyclohexen-1-yl acetate | Menthane monoterpenoids | 9.445 | 195.13867 |
| 91 | NCGC00170014-03!(E)-N-(4-acetamidobutyl)-3-(4-hydroxy-3-methoxyphenyl)prop-2-enamide | Methoxyphenols | 7.675 | 351.15802 |
| 92 | N-Methyl-2-pyrrolidone | N-alkylpyrrolidines | 2.755 | 100.07572 |
| 93 | 2-(2-hydroxybut-3-en-2-yl)-3a,6,6,9a-tetramethyl-2,4,5,5a,7,8,9,9b-octahydro-1H-benzo[e][1]benzofuran-4,5-diol | Naphthofurans | 13.036 | 361.23572 |
| 94 | [9b-hydroxy-6-(hydroxymethyl)-6,9a-dimethyl-3-oxo-1,3a,4,5,5a,7,8,9-octahydrobenzo[e][2]benzofuran-5-yl] benzoate | Naphthofurans | 6.332 | 406.21790 |
| 95 | NCGC00384938-01!4-hydroxy-7-methoxy-2,3,8-trimethyl-3-(4-methylpent-3-enyl)-2H-benzo[g][1]benzofuran-6,9-dione | Naphthofurans | 15.137 | 369.17294 |
| 96 | Deacetoxy(7)-7-Oxokhivorinic Acid | Naphthopyrans | 14.121 | 519.25208 |
| 97 | Koninginin A | Oxepanes | 8.394 | 285.20438 |
| 98 | 2,4,6-Tribromophenol | P-bromophenols | 0.810 | 326.76001 |
| 99 | Citrusin | Phenolic glycosides | 5.803 | 327.14191 |
| 100 | Benzoic acid + 1O, O-Hex | Phenolic glycosides | 3.671 | 299.07755 |
| 101 | Phorbol-12-Myristate-13-Acetate | Phorbol esters | 9.074 | 617.40668 |
| 102 | Phosphatidylethanolamine (20:3/16:1) Abbr: hLPoPE | Phosphatidylethanolamines | 0.768 | 778.44714 |
| 103 | PG 36:3 | Phosphatidylglycerols | 14.156 | 771.51923 |

| | | | | |
|---|---|---|---|---|
| 104 | PG 40:7 | Phosphatidylglycerols | 15.822 | 819.52777 |
| 105 | Benzoic acid + 2O, O-Pen | p-Hydroxybenzoic acid alkyl esters | 3.504 | 285.06158 |
| 106 | 4-Hydroxy-1-(2-hydroxyethyl)-2,2,6,6-tetramethylpiperidine | Piperidines | 1.565 | 202.17987 |
| 107 | 4-Hydroxy-1-(2-hydroxyethyl)-2,2,6,6-tetramethylpiperidine | Piperidines | 0.827 | 202.18007 |
| 108 | Pleiomutinine | Pleiocarpaman alkaloids | 14.124 | 615.36859 |
| 109 | rutamarin | Psoralens | 14.875 | 355.15707 |
| 110 | NCGC00380843-01!5,7-dihydroxy-2-(4-methoxyphenyl)-3-[3,5,7-trihydroxy-2-(4-hydroxyphenyl)-3,4-dihydro-2H-chromen-6-yl]-2,3-dihydrochromen-4-one | Pyranoisoflavonoids | 6.445 | 557.14349 |
| 111 | Trachelanthine | Pyrrolizidines | 0.827 | 302.19620 |
| 112 | Trachelanthine | Pyrrolizidines | 3.086 | 302.19559 |
| 113 | (1-acetyloxy-3-hydroxy-6,8a-dimethyl-7-oxo-3-propan-2-yl-2,3a,4,8-tetrahydro-1H-azulen-4-yl) 4-hydroxybenzoate | Sesquiterpenoids | 14.606 | 429.19373 |
| 114 | NCGC00385008-01!3-[4-[5-(acetyloxymethyl)-3,4-dihydroxyoxolan-2-yl]oxy-3-(2,5-dihydroxy-6-methylhept-6-en-2-yl)-6,9a,9b-trimethyl-7-prop-1-en-2-yl-1,2,3,3a,4,5,5a,7,8,9-decahydrocyclopenta[a]naphthalen-6-yl]propanoic acid | Sesterterpenoids | 4.215 | 682.45111 |
| 115 | NCGC00385201-01_C45H76O19_(2alpha,3beta,5alpha,25S)-26-(beta-D-Glucopyranosyloxy)-2,22-dihydroxyfurostan-3-yl 2-O-(6-deoxy-alpha-L-mannopyranosyl)-beta-D-glucopyranoside | Steroidal saponins | 9.320 | 903.49701 |
| 116 | Timosaponin B II | Steroidal saponins | 9.356 | 921.50745 |
| 117 | DIHYDROMUNDULETONE | Stilbenes | 4.537 | 423.17847 |
| 118 | Azuleno(5,6-c)furan-1(3H)-one, 4,4a,5,6,7,7a,8,9-octahydro-3,4,8-trihydroxy-6,6,8-trimethyl- | Terpene lactones | 13.207 | 283.14972 |
| 119 | NCGC00385670-01!N-(10-hydroxy-10-methylundecyl)acetamide | Tertiary alcohols | 11.941 | 226.21712 |
| 120 | Rolitetracycline | Tetracyclines | 4.861 | 528.23639 |
| 121 | "(3R,5S,7R,8R,9S,10S,12S,13R,14S,17R)-17-((2R,5R)-5,7-dihydroxyheptan-2-yl)-10,13-dimethylhexadecahydro-1H-cyclopenta[a]phenanthrene-3,7,12-triol" | Tetrahydroxy bile acids, alcohols and derivatives | 14.547 | 421.33588 |
| 122 | (2S,3S,4S,5R,6R)-4-{[(2S,3R,4S,5R,6R)-4,5-dihydroxy-6-(hydroxymethyl)-3-{[(2S,3R,4R,5R,6S)-3,4,5-trihydroxy-6-methyloxan-2-yl]oxy}oxan-2-yl]oxy}-3-hydroxy-6-{[(1S,2R,4S,5R,10S,13R,17S)-2-hydroxy-4,5,9,9,13,20,20-heptamethyl-24-oxahexacyclo[15.5.2.0?,??.0?,??.0?,??.0?,??]tetracosan-10-yl]oxy}-5-{[(2S,3R,4S,5S,6R)-3,4,5-trihydroxy-6-(hydroxymethyl)oxan-2-yl]oxy}oxane-2-carboxylic acid | Triterpene saponins | 7.492 | 1127.56128 |
| 123 | NCGC00385302-01_C42H68O14_(3beta,5xi,9xi)-3-{[2-O-(beta-D-Glucopyranosyl)-beta-D-glucopyranosyl]oxy}-23-hydroxyolean-12-en-28-oic acid | Triterpene saponins | 8.552 | 819.45264 |
| 124 | (1S,2R,4aS,6aR,6bR,10S,12aR,14bS)-1,2,6b,9,9,12a-hexamethyl-10-[(2R,3R,4S,5S,6R)-3,4,5-trihydroxy-6-methyloxan-2-yl]oxy-2,3,4,5,6,6a,7,8,8a,10,11,12,13,14b-tetradecahydro-1H-picene-4a,6a-dicarboxylic acid | Triterpene saponins | 8.275 | 633.40009 |
| 125 | Asperosaponin VI | Triterpene saponins | 9.146 | 951.49536 |
| 126 | Notoginsenoside R1 | Triterpene saponins | 7.725 | 955.52350 |
| 127 | Diacetylpyxinol | Triterpene saponins | 4.067 | 1138.86108 |
| 128 | Ecliptasaponin D | Triterpene saponins | 9.103 | 635.41736 |
| 129 | NCGC00385302-01_C42H68O14_(3beta,5xi,9xi)-3-{[2-O-(beta-D-Glucopyranosyl)-beta-D-glucopyranosyl]oxy}-23-hydroxyolean-12-en-28-oic acid | Triterpene saponins | 9.103 | 797.47089 |

| | | | | |
|---|---|---|---|---|
| 130 | Saikosaponin B3 | Triterpene saponins | 14.950 | 813.51215 |
| 131 | Soyasapogenol B base + O-DDMP, O-HexA-HexA | Triterpene saponins | 8.525 | 937.47797 |
| 132 | Soyasapogenol B base + O-HexA-Hex-Pen | Triterpene saponins | 9.117 | 927.49426 |
| 133 | NCGC00385731-01_C46H74O17_(3beta,5xi,9xi,16alpha)-3-{[alpha-L-Arabinopyranosyl-(1->2)-alpha-L-arabinopyranosyl-(1->6)-beta-D-glucopyranosyl]oxy}-16-hydroxyolean-12-en-28-oic acid | Triterpene saponins | 9.361 | 897.48383 |
| 134 | (2S,3S,4S,5R,6R)-6-[[(3S,6aR,6bS,8aR,9R,12aS,14bR)-9-hydroxy-4,4,6a,6b,8a,11,11,14b-octamethyl-1,2,3,4a,5,6,7,8,9,10,12,12a,14,14a-tetradecahydropicen-3-yl]oxy]-5-[(2R,3R,4S,5R,6S)-6-carboxy-4,5-dihydroxy-3-[(2S,3R,4R,5R,6S)-3,4,5-trihydroxy-6-methyloxan-2-yl]oxyoxan-2-yl]oxy-3,4-dihydroxyoxane-2-carboxylic acid | Triterpene saponins | 9.802 | 939.49640 |
| 135 | Madecassoside | Triterpene saponins | 9.117 | 973.49994 |
| 136 | Ginsenoside compound K | Triterpene saponins | 16.069 | 621.43750 |
| 137 | Soyasapogenol B base + O-DDMP, O-HexA-HexA | Triterpene saponins | 8.812 | 935.47809 |
| 138 | Scabioside C | Triterpenoids | 9.812 | 767.46039 |
| 139 | NCGC00386040-01_C52H86O22_alpha-L-Arabinopyranoside, (3beta,5xi,9xi,16alpha)-13,28-epoxy-16,23-dihydroxyoleanan-3-yl O-beta-D-glucopyranosyl-(1->2)-O-[O-beta-D-xylopyranosyl-(1->2)-beta-D-glucopyranosyl-(1->4)]- | Triterpenoids | 6.909 | 1085.54907 |
| 140 | Ursolic acid | Triterpenoids | 8.625 | 457.36865 |
| 141 | betulonaldehyde | Triterpenoids | 8.625 | 439.35764 |
| 142 | MLS000563157-01! | Triterpenoids | 8.972 | 473.36392 |
| 143 | O-Tigloylgymnemagenin, 21- | Triterpenoids | 8.625 | 589.41162 |
| 144 | NCGC00380239-01!16-hydroxy-4,4,8,10,14-pentamethyl-17-(4,5,6-trihydroxy-6-methylheptan-2-yl)-1,2,5,6,7,9,11,12,15,16-decahydrocyclopenta[a]phenanthren-3-one [IIN-based on: CCMSLIB00000848752] | Triterpenoids | 9.103 | 437.34390 |
| 145 | Echinocystic acid | Triterpenoids | 12.601 | 473.36313 |
| 146 | Ganoderic acid LM2 | Triterpenoids | 9.002 | 537.28400 |
| 147 | Wilforlide A | Triterpenoids | 9.103 | 455.35361 |
| 148 | 16beta-Hydroxytrametenolic acid | Triterpenoids | 9.103 | 473.36301 |
| 149 | Scabioside C | Triterpenoids | 9.146 | 767.46033 |
| 150 | Glycyrrhetinic Acid | Triterpenoids | 8.297 | 471.34729 |
| 151 | Saikogenin D | Triterpenoids | 9.795 | 473.36395 |
| 152 | Kalopanaxsaponin H | Triterpenoids | 8.625 | 913.51764 |
| 153 | Ganoderic Acid G | Triterpenoids | 6.082 | 533.30676 |
| 154 | Ginsenoside Rg6 | Triterpenoids | 9.378 | 767.47894 |
| 155 | Ursolic acid | Triterpenoids | 14.206 | 439.35703 |
| 156 | Ursolic Acid | Triterpenoids | 14.156 | 455.35248 |
| 157 | Cauloside C | Triterpenoids | 9.802 | 811.44885 |
| 158 | MPL-dm | Tyrosols and derivatives | 8.071 | 254.17589 |
| 159 | 3-ureidopropionic acid | Ureas | 14.124 | 133.10130 |
| 160 | Ceridimine | Vobasan alkaloids | 10.787 | 497.29071 |
| 161 | Ceridimine | Vobasan alkaloids | 9.002 | 497.29086 |

| | | | | |
|---|---|---|---|---|
| 162 | 2-[1-[1-hydroxy-10,13-dimethyl-3-[3,4,5-trihydroxy-6-[[3,4,5-trihydroxy-6-(hydroxymethyl)oxan-2-yl]oxymethyl]oxan-2-yl]oxy-2,3,4,7,8,9,11,12,14,15,16,17-dodecahydro-1H-cyclopenta[a]phenanthren-17-yl]ethyl]-4,5-dimethyl-2,3-dihydropyran-6-one | Withanolide glycosides and derivatives | 0.768 | 784.44135 |
| 163 | Schweinfurthin F | Xanthenes | 10.787 | 479.27991 |

**Supplementary Table 2.** List of proteins identified in *B. monnieri* -derived nanovesicles.

| N. | ID | Protein name | Mol. Weight [kDa] | Score |
|---|---|---|---|---|
| **Bacopa database (total number =5)** | | | | |
| 1 | A0A6H1YDU0 | DNA-directed RNA polymerase subunit beta | 156.92 | 1.79 |
| 2 | A0A8F2EG55 | Geraniol 10-hydroxylase | 82.945 | 0.22194 |
| 3 | A0A6H1YDW2 | Large ribosomal subunit protein uL2c | 30.041 | 0.19163 |
| 4 | Q5GAB1 | Maturase K | 60.43 | 0.077347 |
| 5 | A0A6H1YDJ2 | Protein Ycf2 | 266.69 | 0.056272 |
| **Plantaginaceae database (total number =27)** | | | | |
| 1 | S5S4I5 | Superoxide dismutase [Cu-Zn], chloroplastic (EC 1.15.1.1) | 15.485 | 78.242 |
| 2 | Q5ZF66 | Superoxide dismutase [Cu-Zn] | 11.658 | 11.22 |
| 3 | A0ABR0CLV4 | Uncharacterized protein | 78.234 | 2.922 |
| 4 | A0ABR0D776 | Uncharacterized protein | 45.935 | 2.7808 |
| 5 | A0ABD3UNT0 | Flavonoid-6-hydroxylase | 59.678 | 2.2708 |
| 6 | A0ABD3SN54 | Probable cytosolic iron-sulfur protein assembly protein CIAO1 homolog | 39.355 | 2.0644 |
| 7 | A0ABD3SZG1 | Uncharacterized protein | 49.444 | 1.9187 |
| 8 | A0ABR0CUH1 | Uncharacterized protein | 73.649 | 1.9187 |
| 9 | A0ABR0DLF1 | Uncharacterized protein | 100.26 | 1.9187 |
| 10 | A0ABD3S4K3 | Uncharacterized protein | 19.59 | 0.6685 |
| 11 | A0ABR0DGP5 | Uncharacterized protein | 66.325 | 0.6685 |
| 12 | A0ABR0CWA1 | Uncharacterized protein | 71.856 | 0.6685 |
| 13 | A0ABR0DVJ3 | Uncharacterized protein | 23.417 | 0.5689 |
| 14 | A0ABR0D689 | Uncharacterized protein | 88.988 | 0.5689 |
| 15 | A0ABR0DR21 | Uncharacterized protein | 72.098 | 0.5689 |
| 16 | W8PFD7 | DIVARICATA (Fragment) | 13.723 | 0.51278 |
| 17 | A0ABD3RSB0 | Uncharacterized protein | 96.69 | 0.41211 |
| 18 | A0ABD3S8X4 | Peptidase A1 domain-containing protein | 48.581 | 0.38042 |
| 19 | A0ABD3S4Y5 | Uncharacterized protein | 63.747 | 0.22477 |
| 20 | A0ABR0CLJ3 | Uncharacterized protein | 128.52 | 0.22477 |
| 21 | A0ABR0DRA9 | Uncharacterized protein | 361.18 | 0.22477 |
| 22 | A0ABR0D801 | Uncharacterized protein | 65.755 | 0.2132 |
| 23 | A0ABR0DQQ7 | Uncharacterized protein | 78.256 | 0.16298 |
| 24 | A0ABR0CZX3 | Uncharacterized protein | 60.654 | 0.14482 |
| 25 | A0ABD3SYJ9 | RING-type E3 ubiquitin transferase (EC 2.3.2.27) | 19.724 | 0.13147 |
| 26 | A0ABD3S179 | Uncharacterized protein | 32.449 | 0.03297 |
| 27 | A0ABD3RUT3 | 2Fe-2S ferredoxin | 19.963 | -0.00089036 |
